# Lineage tracing of newly accrued nuclei in skeletal myofibers uncovers distinct transcripts and interplay between nuclear populations

**DOI:** 10.1101/2023.08.24.554609

**Authors:** Chengyi Sun, Casey O. Swoboda, Michael J. Petrany, Sreeja Parameswaran, Andrew VonHandorf, Matthew T. Weirauch, Christoph Lepper, Douglas P. Millay

## Abstract

Multinucleated skeletal muscle cells have an obligatory need to acquire additional nuclei through fusion with activated skeletal muscle stem cells when responding to both developmental and adaptive growth stimuli. A fundamental question in skeletal muscle biology has been the reason underlying this need for new nuclei in syncytial cells that already harbor hundreds of nuclei. To begin to answer this long-standing question, we utilized nuclear RNA-sequencing approaches and developed a lineage tracing strategy capable of defining the transcriptional state of recently fused nuclei and distinguishing this state from that of pre-existing nuclei. Our findings reveal the presence of conserved markers of newly fused nuclei both during development and after a hypertrophic stimulus in the adult. However, newly fused nuclei also exhibit divergent gene expression that is determined by the myogenic environment to which they fuse. Moreover, accrual of new nuclei through fusion is required for nuclei already resident in adult myofibers to mount a normal transcriptional response to a load-inducing stimulus. We propose a model of mutual regulation in the control of skeletal muscle development and adaptations, where newly fused and pre-existing myonuclear populations influence each other to maintain optimal functional growth.

## Introduction

In vertebrate animals, skeletal muscle is a highly plastic tissue that can adapt its structure and function in response to diverse stimuli. Unlike most other cell types, mature muscle cells (myofibers) possess hundreds of nuclei. This multinucleation is established during embryonic and postnatal development through the fusion of progenitors that have differentiated from muscle stem cells (MuSCs or satellite cells) ^1–3^. Adult muscle can also acquire more nuclei through the activation and fusion of a normally quiescent MuSC population ^4–13^. Since nuclei provide templates for the biosynthetic capacity of cells, the conventional explanation for multinucleation in skeletal muscle is that there needs to be sufficient DNA content to support large cytoplasmic volumes associated with myofibers ^11,14–23^. Indeed, myofibers rank among the largest cells in mammals. In mice, these cells can extend up to 0.5 cm in length and occupy a volume of 7 nL, whereas human myofibers can reach lengths of up to 42 cm, with a volume of approximately 1000 nL ^3,24,25^. To establish size, grow, and adapt, myofibers have two sources of transcriptional templates, either nuclei already present in the syncytium or newly fused nuclei. However, it is unclear how these nuclear sources transcriptionally coordinate muscle growth and adaptations ^14,26,27^.

Despite previous controversy about the requirement of MuSC-mediated myonuclear accretion for muscle growth in the adult, there is now reasonable agreement that pre-existing myonuclei can increase output within limits, but additional nuclei are needed for sustained functional growth ^28–34^. Potential models to explain the need for new myonuclei is that they aid in adaptations by increasing gene dosage or they make unique transcriptional contributions needed for growth that cannot be elicited from pre-existing nuclei. Assessing the role of myonuclear number and accretion in muscle development and growth has proven to be complex perhaps because studies have mainly utilized phenotypic analyses at the level of the whole muscle or individual myofibers. There is minimal information at nucleus-level resolution, but if available, it could help explain from a molecular perspective the need for multinucleation and new nuclei in the adult for adaptations. The development of next-generation sequencing (NGS) strategies such as single nucleus RNA-sequencing (snRNA-seq) could be helpful to parse apart the molecular identities of newly fused and pre-existing myonuclei ^35^. snRNA-seq has been performed on human and mouse muscle, and transcriptional states of distinct myonuclear populations have been detected ^36–40^. However, snRNA-seq approaches lack spatial information and knowledge about the recency of fusion, which are needed to separate contributions of newly fused and pre-existing nuclei. Lineage tracing approaches are also needed to identify recently fused nuclei, but the classical genetic Cre-LoxP system will not work because it causes indelible expression of a marker gene that can diffuse into other nuclei within the shared cytoplasm ^41^.

Here, we report the development of a recombination-independent lineage tracing approach, which we combined with snRNA-seq and bulk nuclear RNA-sequencing (bnRNAseq) strategies to define the transcriptional contributions of newly fused and pre-existing myonuclei to muscle development and adult load-induced adaptations. We found a conserved signature for newly fused myonuclei between development and the adult, which was characterized by elevated expression of *H19* and *Igf2*. During muscle overload in the adult, we observed a distinct transcriptional signature of new myonuclei based on their fusion with previously formed myofibers or fusion to form new myofibers (*de novo*). We also detected a population of pre-existing mature myonuclei that activate a response to overload, associated with expression of *Atf3*, *FlnC,* and *Enah*. We found that fusion and myonuclear accrual are needed to coordinate transcription of pre-existing nuclei during load-induced adaptations and that the syncytial destination of newly fused myonuclei is also a determinant of their transcriptional profiles. Overall, by deploying multiple RNA-seq strategies and a novel mouse model that can distinguish newly fused from pre-existing myonuclei we have identified that these two populations of myonuclei influence each other during developmental and adaptive growth.

## Results

We sought to effectively and reliably determine the transcriptional identity of newly fused myonuclei within a syncytium. To this end, we first tested if a potential newly fused myonuclear population could be detected by snRNAseq data from postnatal (P) day 10, when fusion is ongoing. We previously profiled transcriptomes at this time point from all nuclear populations from tibialis anterior muscles ^39^. Using these data, we selected only myogenic populations including MuSCs, myoblasts, and mature myonuclei to display by Uniform Manifold Approximation and Projection (UMAP) (Figure 1A). MuSCs and myonuclear clusters were identified through distinct expression of canonical marker genes including *Pax7* (MuSCs), *Myog* (myoblasts), *Myh4* (Type IIb myonuclei), *Myh1* (Type IIx myonuclei), *Myh2* (Type IIa myonuclei, and *Myh7* (Type I myonuclei) (Figure 1B). We also identified two myonuclear populations (Dev 1 and Dev 2) that possessed low levels of *Pax7* and *Myog* and also expressed low levels of *Myh isoforms* (Figure 1B), suggesting this population may represent a transcriptional state that has progressed past differentiation but not reached full maturity as myonuclei. PHATE trajectory analysis also inferred that Dev 1 nuclei fall closest to a myonuclear identity in the lineage, while Dev 2 nuclei are further distanced from bona fide myonuclei (Figure 1C). Unbiased analysis of gene expression in these myonuclear clusters revealed that Dev 1 nuclei were enriched for specific genes, while Dev 2 nuclei exhibited a continuum of gene expression that aligns with all myogenic populations (Figure 1D). We then integrated snRNAseq data from P10 muscle with snRNAseq data from mice that were 5 months of age, since there is limited fusion once muscle fully develops. Here, we found an enrichment of only one Dev myonuclear population from P10 muscle (Figure 1E). Comparison of this population to the Dev populations found in Figure 1A showed a similarity to Dev 1 (Figure 1F). The Dev 2 gene signature did not yield a unique cluster after integration, therefore we focused our analysis on Dev 1. Gene expression in these nuclei could be separated into one group (*H19*, *Igf2*, *Dlg2*) that were increased in Dev nuclei compared to MuSCs and decreased in other myonuclear populations at P10 and 5 months of age (Figure 1F, pink bar). In contrast, there is also a group of genes (*Utrn*, *Dpp6*, *Myzap*) expressed in Dev nuclei but at a reduced level compared to MuSCs (Figure 1F, purple bar), suggesting that these genes are more enriched in the differentiation process. Thus, we detected a myonuclear population that correlates with ongoing fusion in muscle suggesting they could represent newly fused nuclei.

**Figure 1.**
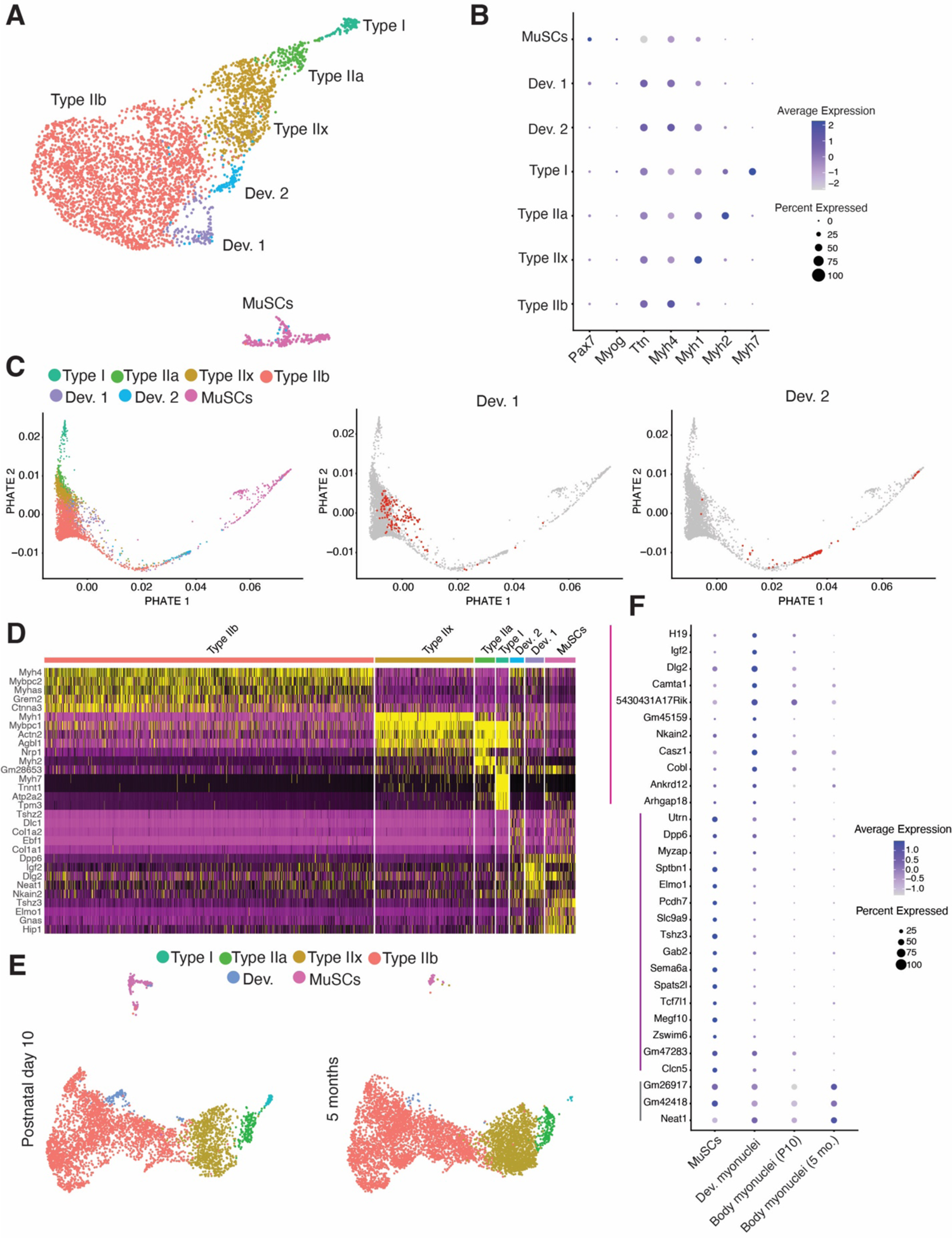
Distinct myonuclear populations are associated with times of fusion. (A) Uniform Manifold Approximation and Projection (UMAP) visualization represents seven color-coded myogenic nuclei clusters identified by snRNA-seq. Data were generated from tibial anterior (TA) muscles at postnatal (P) day 10. (B) Dot plot depicting canonical markers of the various myogenic states. Dot size denotes the percentage of nuclei expressing the gene. (C) PHATE trajectory illustrating myogenic populations presented in (A) (left), with Dev. 1 and Dev. 2 populations highlighted in the same PHATE plot on the right. (D) Heatmap of myogenic nuclei from P10 TA muscle shows a distinct transcriptional signature. (E) Integrated dataset displays myogenic nuclei from TA muscle at P10 and 5 months of age (Split UMAP view). (F) Dot plot demonstrates upregulated genes in Dev. myonuclei. Dot size denotes the percentage of nuclei expressing the gene.

### A recombination-independent tracking system labels recently fused myonuclei

To determine if the Dev myonuclei identified through snRNA-seq truly correspond to newly fused myonuclei we needed to develop a lineage tracing strategy to distinguish if they are present within the syncytium and recently fused. Traditional Cre/LoxP methods are not suitable here as they result in an indelible reporter expression that can label neighboring myonuclei in a syncytium. To overcome these issues, we developed a non-recombination system where the transcription of the nuclear reporter is temporally regulated but produces a long-lived reporter protein. We utilized mice where a reverse tetracycline transactivator (rtTA) construct was knocked-in to the *Pax7* locus (*Pax7*^rtTA^), which results in transcription of rtTA in MuSCs, but rtTA requires Doxycycline (Dox) to bind to tetracycline-responsive elements (TRE) to induce transcription. *Pax7*^rtTA^ mice were bred with mice containing a H2B-GFP expression cassette under control of a TRE (TRE^H2B-GFP^) ^42^. Treatment of *Pax7*^rtTA^; TRE^H2B-GFP^ mice with Dox should activate rtTA in MuSCs leading to expression of H2B-GFP mRNA and protein (Figure 2A). As MuSCs become activated and differentiate, *Pax7* is downregulated resulting in no transcription of rtTA or H2B-GFP thereby reducing mRNA levels of the nuclear reporter prior to fusion, but H2B-GFP protein has a long-half life ^43^ and should be a sustained nuclear mark as the myogenic progenitor fuses into the syncytium (Figure 2A).

**Figure 2.**
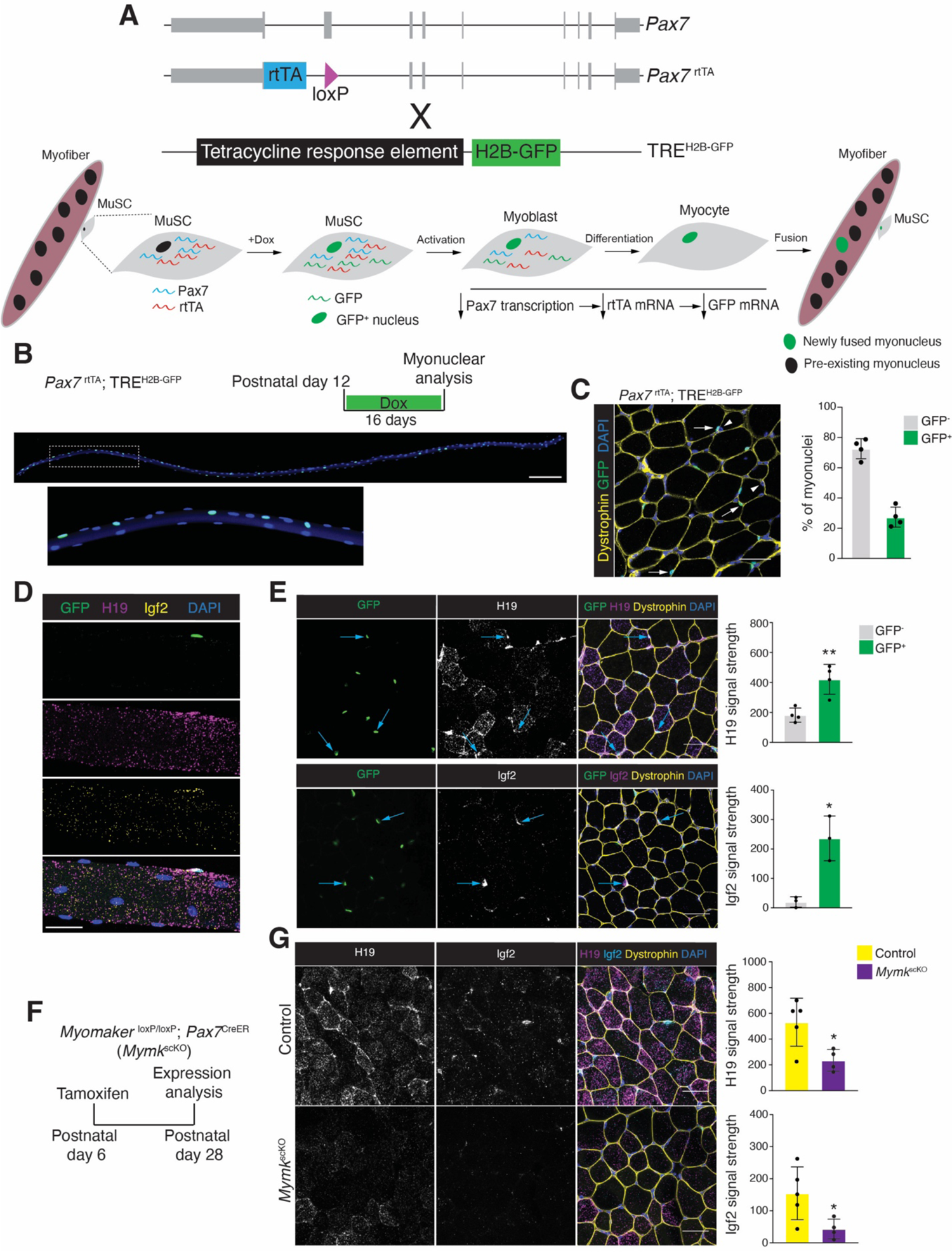
A recombination-independent nuclear tracking system distinguishes newly fused nuclei and allows evaluation of gene expression. (A) Schematic of the *Pax7*^rtTA^; TRE^H2B-GFP^ alleles and Dox-inducible lineage tracing system. (B) Representative images of a single myofiber isolated from the Extensor digitorum longus (EDL) muscle at P28 after 16 days of Dox treatment. The bottom panel represents a magnified field of view of the region indicated by the box in the top panel. Scale bar, 200μm. (C) Representative image showing newly fused GFP^+^ myonuclei (white arrows) and pre-existing GFP^-^ myonuclei (arrowhead) (left). Quantification of the percentages of GFP^+^ myonuclei and GFP^-^ myonuclei (right) in TA muscles. Dystrophin was used to outline myofibers. (n=4). Scale bar, 20μm. (D) Representative images of smRNA-FISH for *H19* and *Igf2* on an EDL myofiber, taken from the same mice used in (C). Scale bar, 20μm. (E) Representative images of smRNA-FISH for *H19* and *Igf2* (left) on *Pax7*^rtTA^; TRE^H2B-GFP^ TA muscle at P28. Quantification of signal in GFP^+^ and GFP^-^ is shown on the right. Blue arrows indicate increased signal in newly fused GFP^+^ myonuclei. (n=3-4). Scale bar, 20μm. (F) Experimental design used to delete Myomaker and myonuclear accretion during development is shown. *Myomaker*^loxP/loxP^; *Pax7*^CreER^ mice were treated with tamoxifen to generate fusion-incompetent mice (*Mymk*^scKO^). (G) Representative images of smRNA-FISH for *H19* and *Igf2* (left) from control and *Mymk*^scKO^ TA muscles. Quantification of *H19* and *Igf2* signal is shown on the right. (n=4-5). Scale bar, 20μm. Data are presented as mean ± SD. Statistical tests used were (E) paired t-test; (G) unpaired t-test; *p<0.05, **p<0.01.

We first assessed if MuSCs are labelled in *Pax7*^rtTA^; TRE^H2B-GFP^ mice by administering Dox chow to 5-month-old mice for 5 days. Flow cytometry analysis for MuSCs (VCAM1^+^ Lin^-^) confirmed that 93% were GFP^+^ (Figure S1A). We also determined that 91% of the GFP^+^ cells were MuSCs (Figure S1B), indicating accurate labeling of MuSCs. We next assessed if this system can label newly fused myonuclei. *Pax7*^rtTA^; TRE^H2B-GFP^ mice were administered Dox for two weeks starting at postnatal day 12 (P12) (Figure 2B) when fusion is ongoing. Isolation of myofibers from the extensor digitorum longus (EDL) muscle indicated the presence of GFP^+^ and GFP^-^ nuclei indicating that H2B-GFP does not move throughout the syncytium (Figure 2B). FACS analysis for PCM1^+^ myonuclei ^44^ from gastrocnemius muscles revealed 21% are GFP^+^ but GFP^+^ myonuclei were absent in TRE^H2B-GFP^ mice (Figure S1C). We also found 25% GFP^+^ myonuclei in the tibialis anterior muscle in cryosections, which were identified as myonuclei based on their presence inside dystrophin^+^ myofibers (Figure 2C). Taken together, we detected that this recombination-independent system labels approximately 25% of myonuclei from P12 to P28, which is consistent with the number of nuclei added to myofibers during this period ^3,45^, indicating that this system can accurately distinguish myonuclei based on their recency of fusion.

### The transcriptional contribution of newly fused myonuclei during development and adult muscle overload

Having established that snRNA-seq data revealed a myonuclear population enriched during development, and the generation of a lineage tracing strategy that marks newly fused myonuclei, we set out to test if the Dev population is indeed newly fused and define the transcriptional contributions of newly fused and pre-existing myonuclei during development. Based on gene expression from the snRNA-seq data at P10, *H19* and *Igf2* are enriched in recently fused myonuclei. We directly tested this idea by performing smRNA-FISH for *H19* and *Igf2* in isolated myofibers and on cryosections from *Pax7*^rtTA^; TRE^H2B-GFP^ mice that were treated with Dox from P12 to P28. We observed an enrichment of *H19* and *Igf2* around GFP^+^ myonuclei in isolated EDL myofibers (Figure 2D). Similarly, on cryosections from the TA we found increased *H19* and *Igf2* signal in myofibers with GFP^+^ myonuclei (Figures 2E). To further confirm that newly fused myonuclei contribute *H19* and *Igf2* to the syncytium, we employed a mouse model that blocks fusion after temporal deletion of Myomaker in MuSCs (*Mymk*^loxP/loxP^; *Pax7*^CreER^ (*Mymk*^scKO^))^46^. Myomaker is deleted upon treatment with tamoxifen and this was performed at P6, and muscles were analyzed at P28 (Figure 2F). As expected, *H19* and *Igf2* levels were reduced in TA myofibers where fusion was blocked at P6 (Figure 2G). These results confirm that the Dev population identified through snRNA-seq at P10 are recently fused myonuclei and those nuclei deliver *H19* and *Igf2*.

To more thoroughly characterize expression in newly fused myonuclei, we performed bulk nuclear RNA-sequencing (bnRNA-seq) on nuclei from *Pax7*^rtTA^; TRE^H2B-GFP^ mice that were treated with Dox from P18 to P28. P28 was chosen for analysis because at this stage developmental growth processes are ongoing and there are recently fused nuclei (P18 to P28) and also myonuclei that are more mature (fused before P18); we thus surmised that this plan could lead to starker differences in gene expression. We separated GFP^+^ (most recently fused) and GFP^-^ (pre-existing) through FACS for PCM1 (from TA and Gastroc pooled together), and then performed independent RNA-sequencing on those populations (Figure 3A). Principal component analysis (PCA) revealed strong separation of GFP^+^ and GFP^-^ myonuclei indicating distinct gene expression signatures (Figure 3B). *H19* and *Igf2* were enriched in GFP^+^ myonuclei, consistent with snRNAseq results. We detected 1877 genes more highly expressed in GFP^+^ myonuclei and 2789 genes more highly expressed in GFP^-^ myonuclei (Figure 3C). In particular, *Gm15668, Gm3669, Cpne4, and Btg4* were significantly enriched in GFP^+^ myonuclei based on a 0.01 adjusted p-value and 2-fold change. In addition, we found *Mymk* enriched in GFP^+^ nuclei, which likely remains from the fusion process and thus highlights the fidelity to distinguish expression in these nuclear populations with our system. *Pcdhga1, Amy2a3, Grem2, Lrrtm3, and Kcnn2* were the top downregulated genes in GFP^+^ myonuclei compared to GFP^-^. Gene ontology analysis based on biological process for genes enriched in GFP^+^ myonuclei suggested an effect on membrane processes involving adhesion, ion transport, and synapse regulation (Figure 3D). In contrast, genes enriched in GFP^-^ myonuclei were associated with mature muscle processes such as muscle contraction, sarcomeres, and mitochondrial metabolism (Figure 3D). Indeed, GFP^-^ myonuclei were enriched for *Maf*, a transcription factor identified as a regulator of the mature muscle program in fast myofibers ^40^. These data indicate that nuclei that enter myofibers during postnatal developmental growth take on a role associated with plasma membrane remodeling and do not possess a strong transcriptional signature for processes associated with muscle growth such as sarcomere organization. Since there is a preponderance of evidence showing that nuclear numbers determine myofiber sizes, it is possible that pre-existing myonuclei are the main transcriptional contributors to increases in myofiber volumes through sarcomerogenesis during early postnatal growth. Newly fused nuclei could either be regulating surface area increases through membrane homeostasis or be transcriptionally unfocused on specific muscle processes because the level of volume growth needed can be driven by the pre-existing population, and newly fused nuclei are recruited for sarcomerogenesis during later maturation and growth, when volumes attained exceed the biosynthetic capability of the pre-existing myonuclear population.

**Figure 3.**
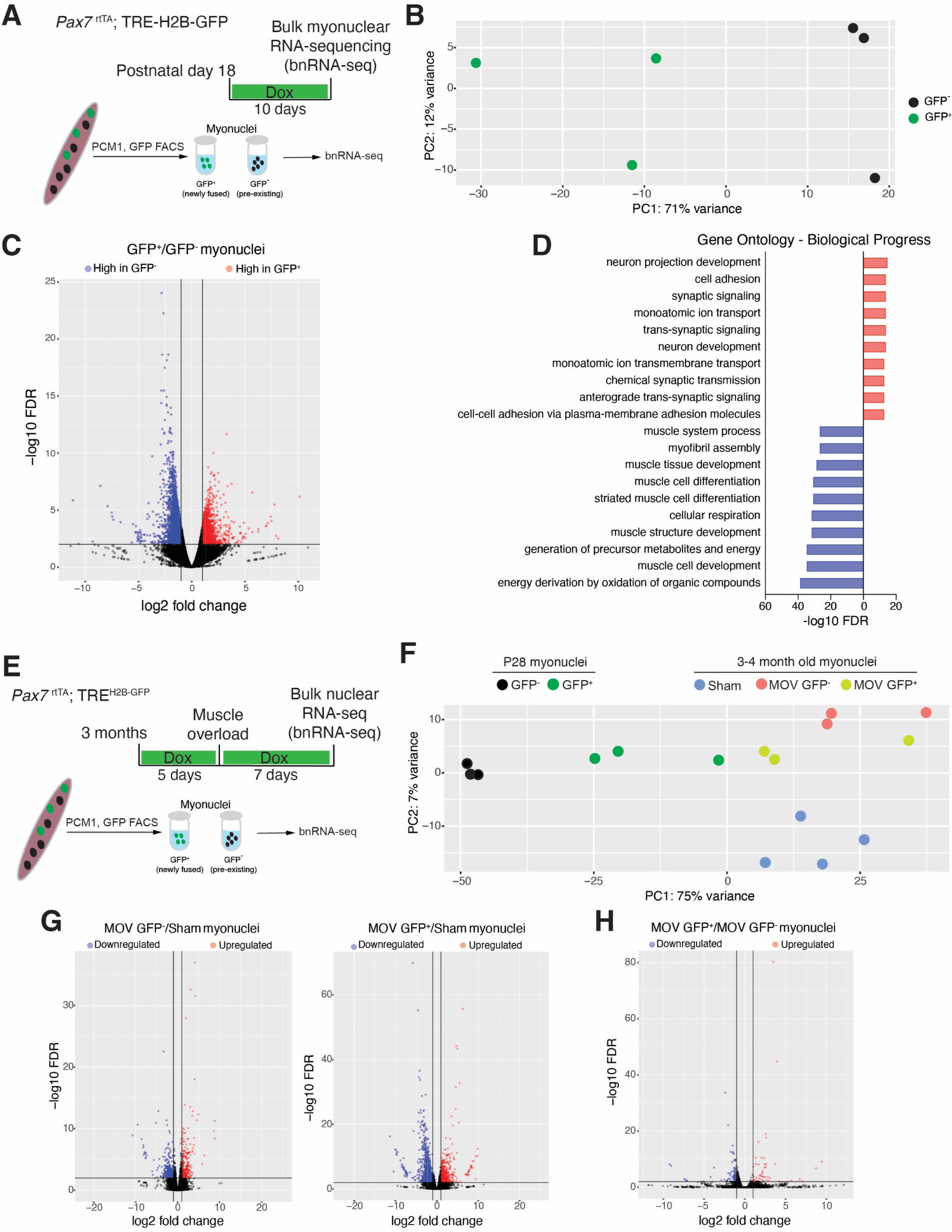
bnRNA-seq analysis comparing newly fused myonuclei acquired during developmental and adult growth. (A) Schematic diagram of the bulk nuclei RNA-seq of GFP^+^ and GFP^-^ myonuclei from *Pax7*^rtTA^; TRE^H2B-GFP^ mice during postnatal development. TA and gastrocnemius muscles at P28 were used after 10 days of doxycycline treatment. (B) Principal component analysis of the top 500 variable genes from bnRNA-seq of newly fused myonuclei (GFP^+^) and pre-existing myonuclei (GFP^-^). (C) Volcano plot depicting DEGs between newly fused myonuclei (GFP^+^) and pre-existing myonuclei (GFP^-^). (D) Gene Ontology (GO) analysis of the DEGs from (C), showing significantly changed biological processes with a false discovery rate (FDR) < 0.01. (E) Schematic diagram for bnRNA-seq of GFP^+^ and GFP^-^ myonuclei from *Pax7*^rtTA^; TRE^H2B-GFP^ mice during adult muscle overload. (F) Principal component analysis of bnRNA-seq data from newly fused myonuclei (GFP^+^) and existing myonuclei (GFP^-^) isolated from plantaris muscle one week after muscle overload, compared with sham myonuclei and developmental myonuclei datasets from (B). (G) Volcano plots reveal up- and down-regulated DEGs in newly fused myonuclei (GFP^+^) or pre-existing myonuclei (GFP^-^) compared with sham myonuclei. (H) Volcano plot reveals up- and down-regulated DEGs from comparison of GFP^+^ and GFP^-^ myonuclei.

To test if the transcriptional signature of newly fused nuclei detected during development is consistent regardless of the growth stimulus that leads to activation of MuSCs and their subsequent fusion with myofibers, we utilized a model of increased load in adult mice through a synergist ablation surgery (muscle overload (MOV)). The gastrocnemius and soleus are removed leaving the plantaris muscle to bear the entire load of the hindlimb resulting in increased load on the plantaris leading to myonuclear accretion and hypertrophy ^47^. We first needed to determine if the *Pax7*^rtTA^; TRE^H2B-GFP^ lineage tracing strategy for newly fused nuclei works in adult muscle. Mice were treated with Dox before and after synergist ablation (muscle overload) (Figure S2A). Analysis of EDL myofibers showed heterogenous GFP expression within myonuclei (Figure S2B). Using FACS to analyze myonuclei (PCM1^+^) we found 25% GFP^+^ myonuclei (Figure S2C). Analysis of GFP^+^ nuclei within dystropin^+^ myofibers also revealed GFP^+^ and GFP^-^ nuclei, where 25-30% were GFP^+^ (Figure S2D). These results are in agreement with the predicted number of new myonuclei that have been observed after muscle overload in previous studies ^31^, highlighting the reliability of *Pax7*^rtTA^; TRE^H2B-GFP^ in tracing newly fused myonuclei during various stimuli.

To assess global gene expression in newly fused and pre-existing myonuclei after adult muscle overload we treated *Pax7*^rtTA^; TRE^H2B-GFP^ mice with Dox five days prior to synergist ablation, then isolated GFP^+^ and GFP^-^ myonuclei one week after surgery (Figure 3E). bnRNA-seq was performed on myonuclei from sham animals, where minimal MuSC activation and myonuclear accrual occurs, and GFP^+^ and GFP^-^ populations from mice that received muscle overload. PCA of these populations showed unique signatures of both GFP^+^ and GFP^-^ myonuclei from muscle overload compared to sham nuclei but we observed a high degree of similarity between the GFP^+^ and GFP^-^ myonuclear transcriptomes (Figure 3F). Comparison of GFP^+^ or GFP^-^ myonuclei to sham myonuclei both reveal hundreds of differentially expressed genes (DEGs) (Figure 3G). Many of these DEGS are shared, with upregulated genes in both GFP^+^ and GFP^-^ myonuclei including *Runx1*, *Ankrd1*, *Ankrd2*, and are consistent with a response to MOV ^48^. We also performed a paired comparison of GFP^+^ and GFP^-^ myonuclei from overloaded muscles, which revealed only 46 upregulated and 78 downregulated DEGs in GFP^+^ myonuclei with FDR<0.01 (Figure 3H), consistent with a high degree of similarity between these nuclear populations. bnRNA-seq only detected a relatively low number of DEGs in newly fused nuclei in MOV, which is in stark contrast to the number of DEGs detected in newly fused nuclei during development, suggesting that newly fused nuclei added during development and MOV are distinct. Indeed, PCA of all of the groups profiled by bnRNA-seq revealed that GFP^+^ nuclei added during development and MOV exhibit a distinct transcriptional profile (Figure 3F). Overall, these data indicate that newly fused and pre-existing myonuclei exhibit distinct transcriptional profiles in certain contexts.

### Divergent contributions of newly fused and pre-existing myonuclei to adult muscle adaptations

While bnRNA-seq did not reveal a robust unique transcriptional signature in GFP^+^ myonuclei after MOV in the adult, the upregulated genes in GFP^+^ myonuclei included *H19*, *Myh3*, *Myh8*, *Tnnt2*, *Tnnt1*. We also found genes associated with extracellular matrix including *Postn*, *Col5a1*, *Col3a1*, *Col12a1*, *Col1a1*, and *Fbn1*. Downregulated genes in GFP^+^ myonuclei (upregulated in GFP^-^ myonuclei) included mature sarcomeric genes such as *Mylk4*, *Myh4*, *Nrap*, and *Xirp2*. We detected some DEGs in GFP^+^ and GFP^-^ myonuclei after MOV, but there is also a high level of variation in gene expression between mice subjected to MOV (Figure 3F), which could be expected given that MOV surgery can lead to a heterogenous response. Due to this variation, we deduced that DEGs could be difficult to identify through a paired analysis of GFP^+^ and GFP^-^ myonuclei, and thus utilized snRNA-seq to determine gene expression in these myonuclear populations after MOV. Here we profiled all myonuclear populations 1 week after synergist ablation using the tet-on HSA^rtTA^; TRE^H2B-GFP^ system ^49^, which results in GFP-labeling of all myonuclei, including newly fused and pre-existing populations (Figure 4A). This system was chosen over the *Pax7*^rtTA^; TRE^H2B-GFP^ system to maximize the number of nuclei obtained for snRNA-seq.

**Figure 4.**
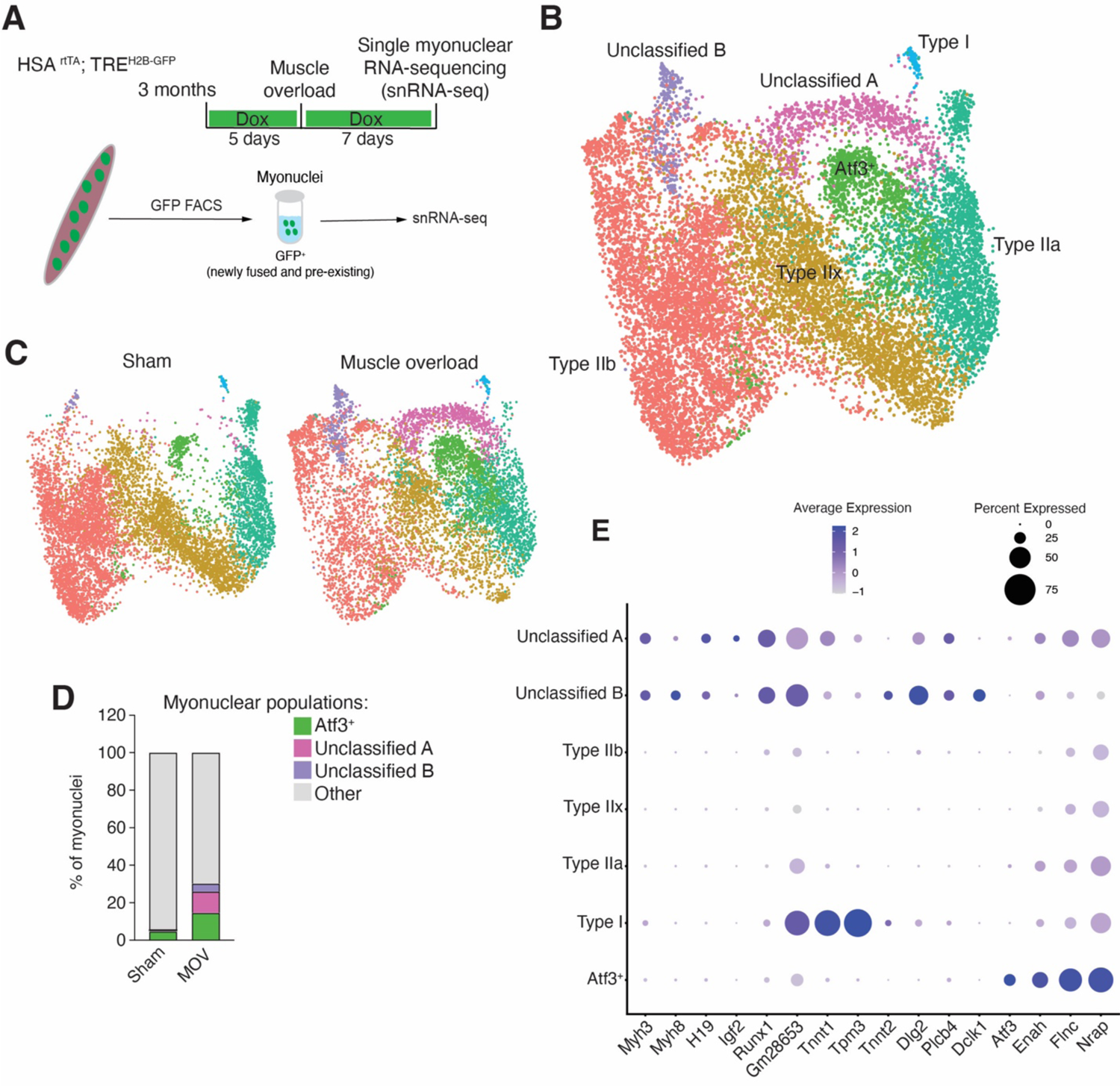
Differential contributions of newly fused and pre-_existing myonuclei to adult_ muscle adaptations. (A) Schematic diagram of the single-nuclei RNA-seq methodology applied to HSA^rtTA^; TRE^H2B-GFP^ mice, where all myonuclei were labeled with GFP and sorted for subsequent sequencing analysis. (B) Integrated UMAP from plantaris muscle following overload stimulation (n=2) or under sham condition (n=1). (C) Split UMAP view of myogenic nuclei shown in (B). (D) Proportion of unclassified A, B, and *Atf3*^+^ nuclear populations. (E) Dot plot depicts selected genes differentially expressed in various myonuclear populations. Dot size represents the percentage of nuclei expressing the gene.

We profiled 11,416 transcriptomes from libraries generated from overloaded plantaris muscle, and 11,182 transcriptomes from libraries generated from the muscle in sham control group. To make sure that the common artifacts were dealt with appropriately in droplet-based transcriptome analysis, we utilized CellBender for removing ambient RNA and Solo for removing doublets from individual samples before integrating the data with Seurat’s SCTransform integration workflow. Sham and MOV data were integrated and myonuclear populations that are in specialized compartments including NMJ, MTJ, and MTJ-B populations were removed ^37–39^. We retained myonuclei not in specialized compartments and performed an additional dimensionality reduction and annotated nuclear clusters based on fiber type (myosin isoform expression) or distinct transcriptional signature (Figure 4B). Four of these seven clusters, belonging to different myofiber types, were characterized by the expression of canonical markers (type I, *Myh7*; type IIa, *Myh2*; type IIx, *Myh1*; type IIb, *Myh4*) and present in both sham and overloaded muscles (Figure S3A). We also detected two populations only present in overload samples and one population that was expressed in sham but enriched in plantaris muscles after overload (Figure 4C). Analysis of the percentage of nuclei confirmed that the emerging populations, referred to as the unclassified A and B, were mainly present in overloaded muscles, while the enriched cluster, named *Atf3*^+^ myonuclei, were present in sham and increased after overload (Figure 4D). Distinct gene expression was detected in the unclassified A and B populations, including *H19*, *Igf2, Runx1,* and embryonic myosin isoforms (*Myh3* and *Myh8*) (Figure 4E and Figure S3B). We also detected differences between unclassified A and B populations, where *Tnnt2* and *Dclk1* are more enriched in unclassified B nuclei (Figure 4E).The Atf3 population was not enriched for *H19*, *Igf2*, *Myh3*, or *Myh8* but instead was enriched for *Atf3*, *Flnc*, *Enah*, and *Ankrd1* (Figures 4E, S3B, and S3C), indicating a similar gene signature to that of *Atf3*^+^ sarcomere assembly myonuclei observed during postnatal muscle growth ^39,50^.

We next utilized the *Pax7*^rtTA^; TRE^H2B-GFP^ system to test if the overload responsive nuclear populations (unclassified A and B, and *Atf3*^+^ myonuclei) were derived from newly fused or pre-existing nuclei. *Pax7*^rtTA^; TRE^H2B-GFP^ mice received a 5-day doxycycline regimen prior to synergist ablation surgery and were continuously treated with doxycycline until sacrifice one week after the procedure (Figure 5A). Analysis of *H19*, *Ig2, Tnnt1, Tnnt2,* and *Atf3* transcripts was performed using smRNA-FISH on cross-sections of plantaris muscle with or without synergist ablation. Our results showed an enrichment of *H19*, *Igf2*, *Tnnt1*, *Tnnt2* surrounding GFP^+^ myonuclei (Figure 5A). In contrast, *Atf3* was enriched in GFP^-^ myonuclei (Figure 5A). Genetic blockade of fusion in *Mymk*^scKO^ mice reduced levels of *H19*, *Igf2*, *Tnnt1* and *Tnnt2*, but not *Atf3*, in myofibers (Figure 5B). These data highlight that unclassified A and B populations in muscle overload are recently fused, and the Atf3 population originates from pre-existing nuclei.

**Figure 5.**
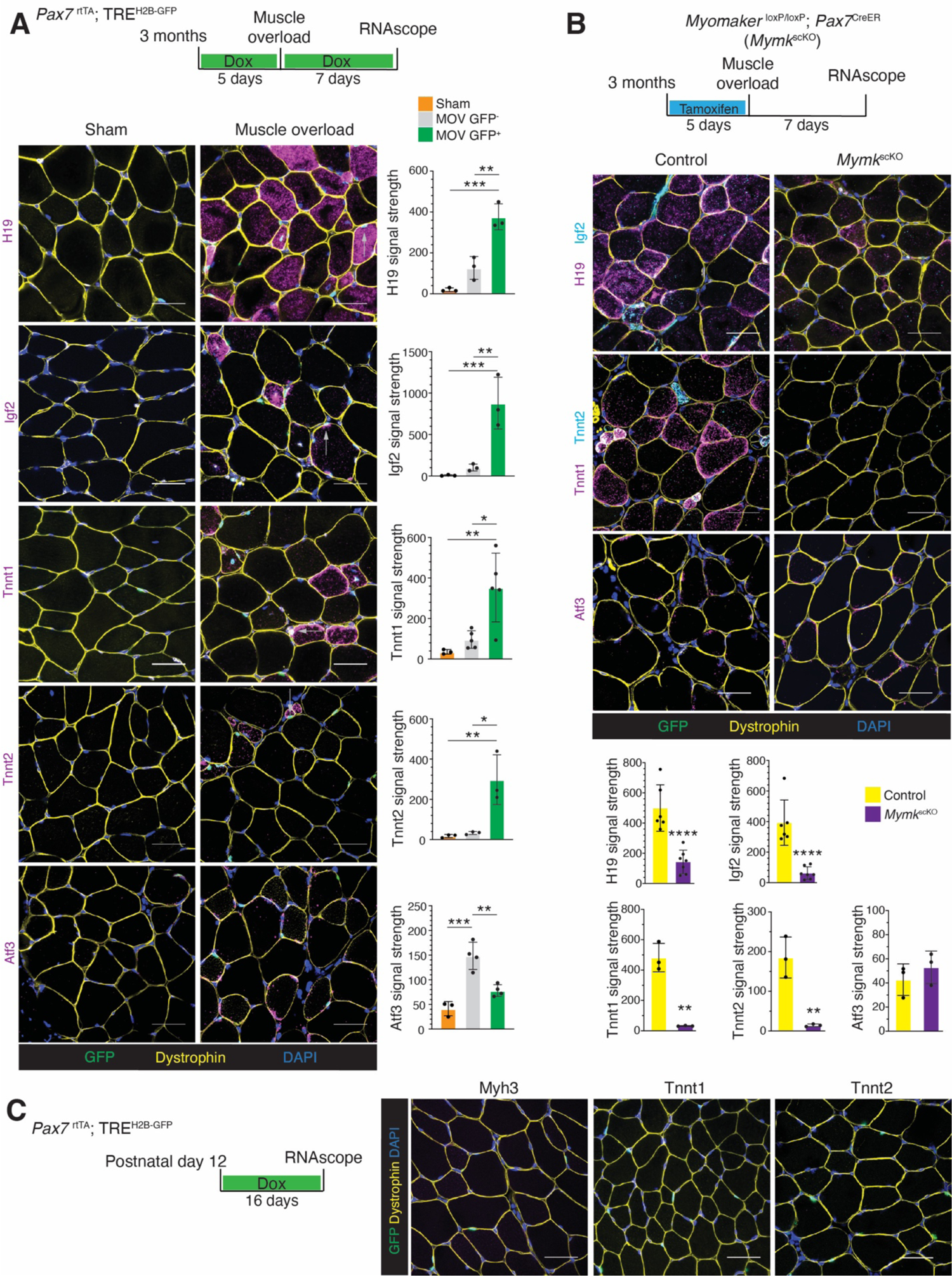
smRNA-FISH analyses validates gene expression in newly fused and pre-existing myonuclei. (A) Representative images of smRNA-FISH on sections from *Pax7*^rtTA^; TRE^H2B-GFP^ muscle for selected genes enriched in newly fused (*H19*, *Igf2*, *Tnnt1*, *Tnnt2*) or pre-existing myonuclei (*Atf3*). Quantification of the signal for each gene is shown on the right. (n=3-5). Scale bar, 20μm. (B) Representative images of smRNA-FISH for the indicated genes on sections from control or *Mymk*^scKO^ mice subjected to muscle overload. Quantification of the signals are shown on the bottom. (n=3-6). Scale bar, 20μm. (C) Representative images of smRNA-FISH for *Myh3*, *Tnnt1*, and *Tnnt2* on sections from *Pax7*^rtTA^; TRE^H2B-GFP^ muscle during postnatal development. (n=3). Scale bar, 20μm. Data are presented as mean ± SD. Statistical tests used were (A) one-way ANOVA with Tukey’s correction for multiple comparisons; (B) unpaired t-test; *p<0.05, **p<0.01, ***p<0.001, ****p<0.0001.

Our analyses of newly fused nuclei during development and after muscle overload suggest that *H19* and *Igf2* are two conserved markers whereas *Tnnt1* (marker of unclassified A and B), *Tnnt2* (marker for unclassified B) and *Myh3* (marker for both unclassified A and B) are unique for muscle overload. smRNA-FISH for *Tnnt1*, *Tnnt2* and *Myh3* on muscle sections from *Pax7*^rtTA^; TRE^H2B-GFP^ mice where newly fused myonuclei were labeled from P12 to P28 revealed no expression (Figure 5C), confirming that these genes are specific markers of newly fused nuclei during MOV.

### The syncytial muscle environment is a determinant of gene expression in newly fused myonuclei

Multiple observations described thus far suggest newly fused myonuclei exhibit differential gene expression. First, PCA of bnRNA-seq data of myonuclear populations during development and MOV revealed separation of newly fused nuclei based on condition (Figure 3F). Second, two newly fused nuclear populations (unclassified A and B) were detected by snRNA-seq after MOV. Third, *Tnnt1*, *Tnnt2*, and *Myh3* were expressed in newly fused nuclei during MOV but not during development (Figure 5C). We next sought to uncover how this divergent gene expression of newly fused nuclei is controlled, which could include that the stimulus directly entrains gene expression in myogenic progenitors prior to fusion or that the syncytial environment to which the new nucleus is fusing into could impact its transcriptional trajectory. To parse apart these possibilities we took advantage of the diversity of cell fusion events during muscle overload, where myoblast-myoblast fusions generate *de novo* myofibers and myoblast-myofiber fusions result in addition of new nuclei to established myofibers ^31,34,51^. In this situation, myogenic progenitors are generally subjected to the same stimulus (increased load) but are fusing into different syncytial environments. Recent evidence indicates that fusion to form *de novo* myofibers is rapid and thus the nuclei are fusing into an immature syncytium ^5^. In contrast, during myoblast-myofiber fusions myogenic progenitors enter an established syncytium governed by mature myonuclei.

The presence of nuclei in the center of myofibers is a hallmark characteristic of *de novo* myofibers and repair of pre-existing myofibers. However, a long-standing question is whether newly fused myonuclei can be deposited in the periphery of myofibers. The data presented in Figure 5 highlight that new nuclei added to myofibers after MOV can be peripherally located. While we observed mainly central GFP^+^ myonuclei in small myofibers (likely *de novo*), we found central and peripheral GFP^+^ myonuclei in larger myofibers (likely pre-existing). This model also allowed us to test if *Myh3*, a classical marker of *de novo* myofibers is also present in larger myofibers that acquired a new nucleus ^52^. We detected *Myh3* mRNA in newly fused myonuclei regardless of whether they are in a central or peripheral location (Figure 6A and 6B). While *Myh3* is not a distinguishing marker of central or peripheral newly fused nuclei after muscle overload, we observed distinctions in *Tnnt2* expression patterns, indicating a divergent transcriptional signature. *Tnnt2* expression, a marker of the unclassified B population, was restricted to smaller, likely *de novo* myofibers after muscle overload (Figure 5B). In contrast, *Tnnt1*, a marker of the unclassified A population, was present in both smaller and larger myofibers (Figure 5B), similar to *Myh3*. These data suggest that the unclassified B population represent nuclei from *de novo* myofibers, which is further supported by data that *Tnnt2* is enriched in embryonic myonuclei compared to postnatal myonuclei ^40^. To assess gene expression alterations in newly acquired myonuclei in *de novo* myofibers (unclassified B) and pre-existing myofibers (unclassified A) we directly compared their transcriptomes to mature myonuclei that are responding to MOV. We found that unclassified B has a more distinct transcriptional signature while unclassified A exhibits more similarity to other myonuclei, especially Type IIa and *Atf3*^+^ myonuclei (Figure 6C). Unclassified B myonuclei seem to share a gene profile with Type IIb myonuclei. Unclassified A and B myonuclei are subjected to the same MOV stimulus as progenitors and yet exhibit distinct gene expression patterns, suggesting that the stimulus itself does not solely explain differential gene expression of newly fused myonuclei in this context, and one of the determinants is the syncytial environment itself.

**Figure 6.**
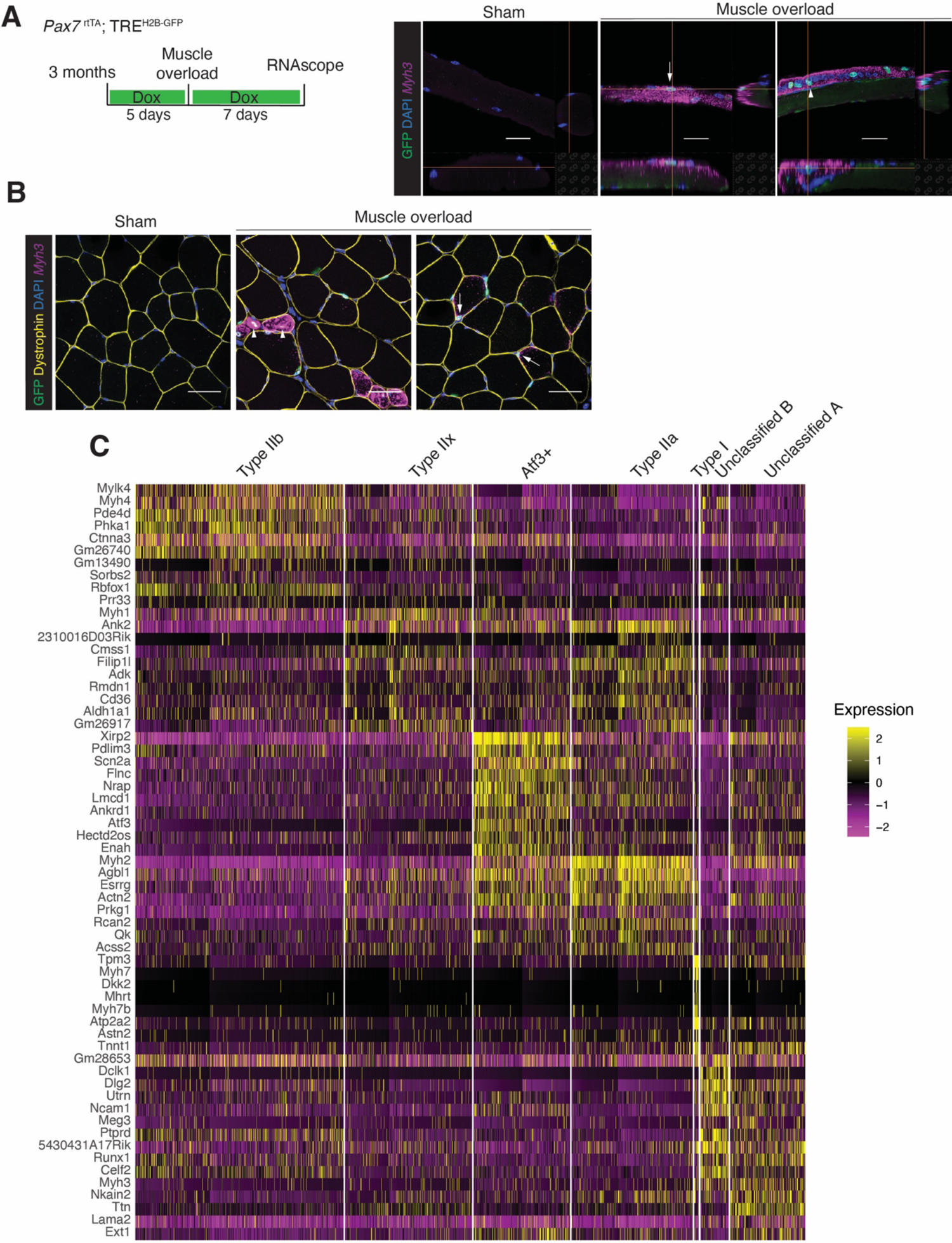
Comparative gene features of newly fused myonuclei in *de novo* and pre-ng myofibers. (A) Representative images of smRNA-FISH for *Myh3* on plantaris myofibers from *Pax7*^rtTA^; TRE^H2B-GFP^ mice undergoing overload stimulation or under sham conditions. *Myh3* transcription was detected in both peripheral (white arrow) and central (arrowhead) newly fused myonuclei. (n=3-4). Scale bar, 20μm. (B) Representative images of smRNA-FISH for *Myh3*, performed on the same mice used in (A), show that both central newly fused GFP^+^ myonuclei (arrowhead) and peripheral newly fused GFP^+^ (white arrow) are enriched for *Myh3* transcription. (n=3-4). Scale bar, 20μm. (C) Heatmap of genes showing distinct transcriptional signature in nuclei of myogenic populations from plantaris muscle subjected to one week of overload stimulation, as used in Figure 4C (n=2).

### Cellular fusion coordinates gene expression of myonuclei after muscle overload

The response of myofibers to muscle overload involves activation of the expression of *Atf3*, *Flnc*, and *Enah* in resident myonuclei, accretion of additional nuclei that express *Tnnt1* into pre-existing myofibers, and generation of *de novo* myofibers whose nuclei express *Tnnt2*. Given this combinatorial response of newly fused and pre-existing myonuclei, that myonuclear accretion is required for proper adaptations of muscle to increased load, and an apparent role of the syncytium in impacting gene expression in newly fused nuclei described here, we asked if the myonuclear accretion process globally impacted gene expression in pre-existing nuclei during adult muscle overload.

We tested this concept by assessing the response of pre-existing myonuclei to muscle overload in the absence of myonuclear accretion by performing bnRNA-seq on myonuclei from WT and *Mymk*^scKO^ mice after overload (Figure 7A). PCA revealed that the transcriptional signature in *Mymk*^scKO^ myonuclei were distinguished from both sham and WT MOV GFP^+^/GFP^-^ myonuclei (Figure 7B). We started by comparing gene expression between the WT MOV myonuclei (GFP^+^ and GFP^-^ combined) and WT sham myonuclei, with the aim to define the transcriptional response to muscle overload in the presence of myonuclear accretion. This analysis identified 587 upregulated genes (Figure S4A). We also contrasted the gene expression of WT MOV myonuclei (GFP^+^ and GFP^-^ combined) with that of *Mymk*^scKO^ MOV myonuclei to identify the response to MOV when myofibers are unable to accrue additional nuclei, which revealed 1,828 upregulated genes (Figure S4B).

**Figure 7.**
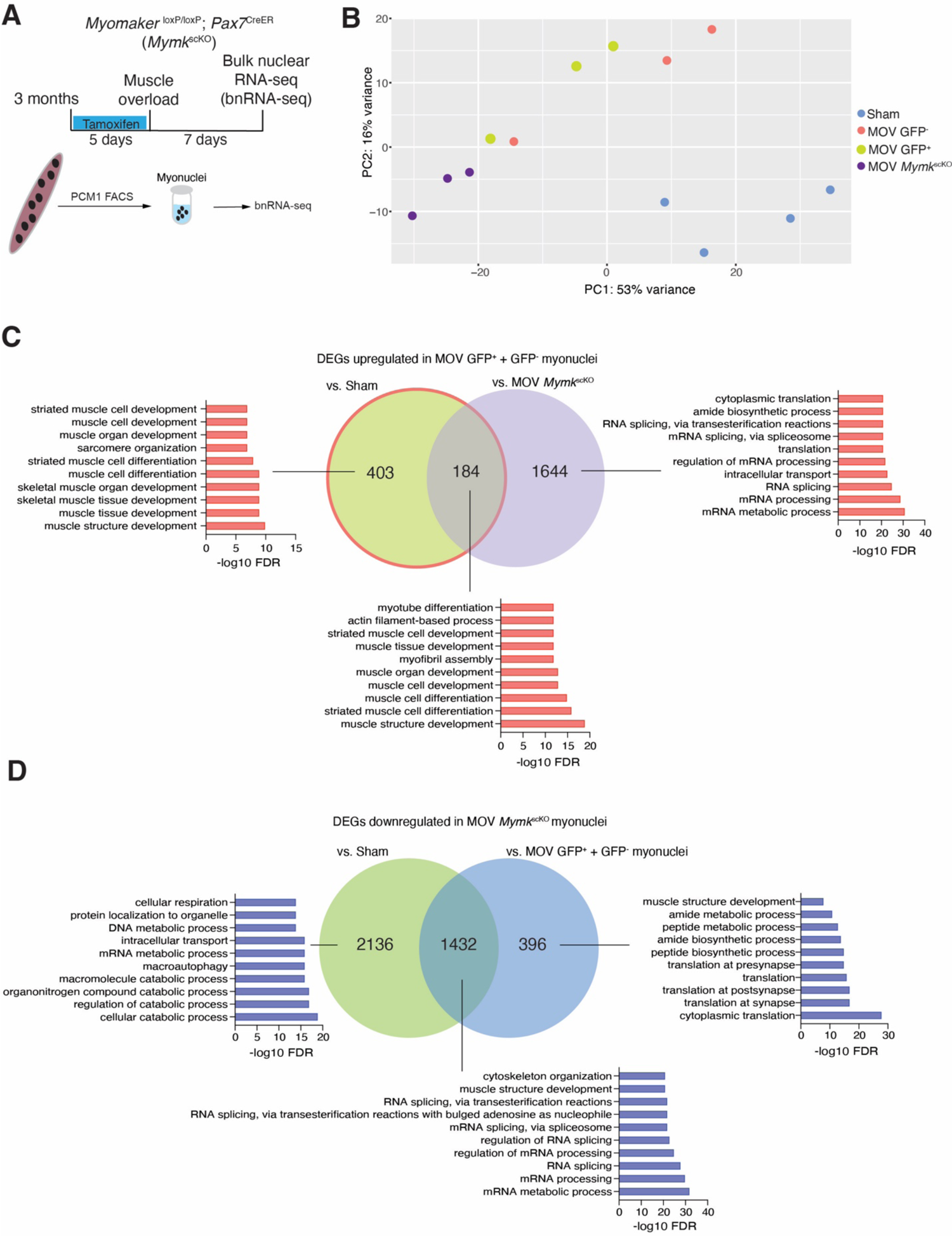
Fusion and myonuclear accrual is needed for pre-existing myonuclei to transcriptionally respond to muscle overload. (A) Schematic diagram of the bulk nuclei RNA-seq methodology for analyzing pre-existing myonuclei adapting to muscle overload (MOV) in the absence of myonuclear accretion. *Myomaker*^loxP/loxP^; *Pax7*^CreER^ mice were treated with tamoxifen to generate fusion-incompetent mice (*Mymk*^scKO^). (B) Principal component analysis of bnRNA-seq from *Mymk*^scKO^ myonuclei isolated from plantaris muscle one week after muscle overload, myonuclei (GFP^+^ and GFP^-^) derived from control mice that underwent the same surgery, and myonuclei from sham mice. (C) Venn diagram showing the distinct or overlapping genes upregulated in WT MOV (GFP^+^ and GFP^-^ combined), compared with WT sham or *Mymk*^scKO^ MOV myonuclei. Gene Ontology analysis is also shown. (D) Venn diagram showing the distinct or overlapping genes downregulated in *Mymk*^scKO^ MOV myonuclei, compared with WT sham or WT MOV (GFP^+^ and GFP^-^ combined) myonuclei. Gene Ontology analysis is also shown.

Of the 587 upregulated DEGs observed in the WT MOV/Sham comparison, 403 were not detected as a DEG in the WT MOV/*Mymk*^scKO^ MOV comparison because they are also activated in *Mymk*^scKO^ muscle (Figure 7C). We consider these 403 genes to represent the transcriptional response to muscle overload that is not dependent on the fusion process. Consistent with our snRNA-seq analysis, we found *Atf3*, *Flnc*, and *Klhl41* within this list of genes, further validating the presence of a transcriptional response elicited by increased load in the resident myonuclear population independent of addition of new nuclei. We determined that 184 DEGs are shared between the WT MOV/Sham and WT MOV/*Mymk*^scKO^ MOV comparisons (Figure 7C). These genes are normally activated in physiological muscle overload adaptations but fail to be expressed when new myonuclei cannot be properly incorporated into the syncytium. These genes, including *Tpm1*, *Tpm2*, *Tpm3*, *Tnnt2*, and *Atp2a2*, are involved in muscle development and contraction regulation, as shown by gene ontology analysis (Figure 7C). These genes also serve as markers for the ‘unclassified A’ and ‘unclassified B’ populations that emerge after MOV, as observed in our snRNA-seq data. The GO analysis of the 403 fusion-independent genes and 184 fusion-dependent genes revealed similar processes associated with muscle development and contraction (Figure 7C), indicating a collaborative mechanism where distinct genes contributed from different myonuclear populations converge on cellular pathways that are needed for proper muscle adaptations.

Additionally, the comparison of gene expression between WT MOV myonuclei and *Mymk*^scKO^ MOV myonuclei revealed 1644 upregulated genes. These genes were not upregulated in the WT MOV/sham comparison suggesting they may represent a discordant transcriptional response from MOV myonuclei in the absence of fusion. Indeed, GO analysis of these genes indicated a role in mRNA processing and protein translation (Figure 7C). Comparison of myonuclear gene expression in *Mymk*^scKO^ MOV to sham found 3568 downregulated DEGs (Figure S4C), indicating that discordant gene expression in the absence of fusion during MOV is due to arrested normal gene expression. Of these 3568 downregulated genes, 1432 are shared with downregulated DEGs with *Mymk*^scKO^ MOV/WT MOV (Figure 7D).

We then analyzed downregulated DEGs in MOV *Mymk*^scKO^/Sham and MOV *Mymk*^scKO^/MOV WT to better characterize the discordant gene expression in the absence of fusion. There were 1432 shared downregulated DEGs in *Mymk*^scKO^ myonuclei irrespective of whether the muscle has been subjected to MOV (Figure 7D). We also identified 2136 genes that uniquely fail to be expressed in *Mymk*^scKO^ when compared with WT Sham, and 396 DEGs only present when the muscle undergoes synergistic ablation (Figure 7D). Besides the impact on mRNA processing, GO analysis revealed that blocking the addition of myonuclei diminishes cytoplasmic translation and ribosomal subunit assembly when muscle receives overload stimulation (Figure 7D). Our findings align with the previously undetected skeletal muscle hypertrophy in *Mymk*^scKO^ MOV mice when the rate of muscle protein synthesis was significantly reduced ^31^. Overall, these data indicate that in the absence of myonuclear accretion, myofibers fail to activate pathways needed for adaptations to muscle overload. Thus, the fusion process not only contributes unique transcripts it also aids in the coordination of gene expression from pre-existing myonuclei in the syncytium.

## Discussion

Although there are multiple syncytial cell types, skeletal myofibers are unique in that they are long-lived cells that not only accrue hundreds of nuclei during development but also can add new nuclei in the adult in response to various stimuli. It has been established that the number of nuclei in skeletal myofibers are a major factor for size and function, and adult myofibers require new nuclei for adaptations ^29–33,53^, which has led to questions about how new nuclei and pre-existing nuclei synergize to maintain a functional syncytium. The main rationale for this study was to determine if newly fused nuclei are transcriptionally distinct from the pre-existing myonuclei. If so, it would suggest that newly fused nuclei possess a transcriptional machinery independent from what is encountered as part of the syncytium. If newly fused nuclei were similar to pre-existing nuclei, it would suggest that they are entrained to become a functioning part of the syncytium. Our results reveal that transcriptional profiles of newly fused nuclei reflect a confluence of both independence and syncytial assimilative forces, both of which are relatively under-discussed aspects of skeletal muscle biology but could represent nodal processes for the regulation of skeletal muscle growth and function.

In this study we overcome a major obstacle in understanding gene expression in myonuclear populations through the development of a lineage tracing model in the mouse that distinguishes newly fused and pre-existing myonuclei. We identify conserved markers of newly fused myonuclei, including *H19* and *Igf2*, which are present in this population during both developmental and adult accrual. Our work also establishes that newly fused myonuclei added to the syncytium during development are distinct from those added during adult load-induced muscle adaptations. We propose a model where the transcriptional trajectory of newly fused nuclei are impacted by the syncytial environment, which is controlled in part by the pre-existing nuclei. Reciprocally, myonuclear accretion is needed for pre-existing myonuclei to appropriately respond to an overload stimulus, indicating a bilateral interaction mechanism between newly fused and pre-existing myonuclei. In addition to defining gene expression of newly fused and pre-existing myonuclei in response to various stimuli, this work establishes an interaction between these nuclear populations in the myofiber syncytium.

What is the purpose of myofibers adding new nuclei during development and adult adaptations? The consequences of cellular fusion in these two scenarios should be considered separately since the physiological and functional activities of myofibers are different and our results show that new nuclei are molecularly distinct. In a previously reported model where fusion is blocked early during postnatal development, remaining myonuclei are able to transcriptionally compensate and support growth ^45^. In contrast, during adult muscle hypertrophy, a fusion blockage results in a lack of growth, indicating adult resident myonuclei are unable to fully compensate ^31–33^. When asking about unique transcriptional contributions of a newly fused myonucleus to a syncytia, one must first determine the bioavailability of certain factors already present within that syncytium. During development, we found that newly fused nuclei are more transcriptionally distinct compared to other nuclei within the syncytium, whereas after muscle overload new nuclei are more similar to pre-existing myonuclei.

For adult muscle adaptations, one idea is that newly fused nuclei are highly specialized and deliver specific transcripts needed for a particular aspect of the adaptation. We confirmed that myonuclear accretion is essential for muscle adaptations in the adult, but our results did not yield an obvious single factor contributed by newly fused nuclei that could explain their necessity for growth. *H19* and *Igf2* are candidate factors since they have known in roles in myogenesis and skeletal muscle ^54–58^, but their function in myofibers after fusion remains to be determined. An alternative interpretation is that the main contribution of new nuclei is to increase the number of DNA copies rather than to contribute unique transcripts that perform a specific function ^59^. However, it is possible that a transcript that currently lacks functional annotation detected in newly fused nuclei does perform a critical function. While new nuclei are providing unique transcripts to the syncytium, these transcripts may not be needed for a specific response to the stimulus but instead coordinate a synergistic response to the stimulus by impacting pre-existing nuclei. Support for the concept that there is global sensing of cellular fusion in myofibers during adult muscle adaptations includes data presented here where pre-existing nuclei transcriptionally decompensate to levels below that of non-stimulated adult myonuclei. These data are consistent with a report that pre-existing myonuclei exhibit divergent gene expression after depletion of MuSCs in response to an 8-week load stimulus ^28^, and further clarify that fusion and myonuclear accrual is the cellular process that pre-existing myonuclei need to mount a functional response to increased load. Our work here highlights the presence of a global sensing mechanism in myofibers where pre-existing myonuclei transcriptionally respond to the accrual of new nuclei, which could include direct communication between the nuclei or indirectly sensing alterations in the syncytium due to lack of new nuclei.

One unanticipated conclusion from our study, which in retrospect should have been obvious given the strong reprogramming abilities of myonuclei ^60,61^, is extent to which the syncytial environment impacts the trajectory of newly fused nuclei. While myogenic progenitors may possess intrinsic differences based on age of the animal, muscle type, and presence of external stimuli ^30,62,63^, leading to heterogenous transcriptional states that could impact the molecular trajectory once it enters myofibers, our data highlight a role for the syncytium as well. The first observation indicating that the syncytial environment impacts newly fused nuclei is that the transcriptional distinctiveness of newly fused nuclei compared to other nuclei within the syncytium was most evident during development. Since there is less of a requirement for new nuclei for muscle growth at this stage given that pre-existing nuclei exhibit an ability to increase output to support larger cytoplasmic volumes, we suspect that the transcriptional differences are due to the overall state of the syncytium during development. Indeed, myonuclei are in a more immature state in the postnatal stage ^40^, where there is likely not a sufficient quantity of mature muscle transcripts to reprogram newly added myonuclei. Another observation supporting a role for the syncytium is that after MOV we detected transcriptional differences in myonuclei that were added to already formed myofibers compared to myonuclei in *de novo* myofibers. After MOV, newly fused myonuclei in *de novo* myofibers, which have a recently established syncytium, were enriched for *Tnnt2* and *Dclk1* transcripts that also mark embryonic myonuclei ^40^. In contrast, newly fused myonuclei in pre-existing myofibers after MOV exhibited transcriptomes similar to those myonuclei that were already present in the syncytium prior to MOV, further suggesting a role for the syncytial environment in regulating the trajectory of newly fused nuclei. It should be noted that the stimulus itself could entrain gene expression of myogenic progenitors prior to fusion, and we are not excluding that as a regulatory mechanism. However, the presence of divergent transcription observed in nuclei added to pre-existing myofibers after MOV compared to those in new myofibers implicates the syncytial environment since these progenitors were subjected to the same load increases. Regardless of the molecular details, we propose that the syncytial environment is one determinant of gene expression in newly fused myonuclei.

In summary, a unique and exclusive function of newly fused myonuclei remains to be determined, but the availability of the lineage tracing model developed here and the subsequent identification of transcriptional markers of newly fused nuclei opens up the possibility to more deeply explore this issue. Our results unexpectedly reveal that newly fused and pre-existing myonuclei reciprocally influence each other to establish the myofiber syncytium and respond to adaptations in adult muscle.

### Limitations of the study

A caveat with our study is that it is difficult to know if detected transcripts in newly fused nuclei are simply a result of the differentiation and fusion process, and we did not analyze gene expression in myogenic progenitors prior to fusion. We analyze gastrocnemius and TA in development and plantaris during MOV, and thus some differences we detect could be due to specific characteristics of those muscles. However, throughout our study we have investigated gene expression in newly fused nuclei using various approaches (RNA-seq and smRNA-FISH) and observed a consistent signature regardless of the source of muscle. For instance, *H19* and *Igf2* were detected as enriched in the following experiments: snRNA-seq at P10 from the TA, bnRNA-seq from GAS/TA at P28, and smRNA-FISH at P28 on isolated EDL myofibers. Given the consistency of DEGs independent of technique, we are confident that our results reliably distinguish gene expression between newly fused and pre-existing myonuclei. From an RNA sequencing perspective, bnRNA-seq is a relatively new approach and we did detect higher than average amount of reads mapping to intergenic regions in all datasets, the relevance of which is unclear, and their presence could limit our capacity to detect more robust transcriptional differences. Finally, it should also be noted that the detection of transcripts using nuclear sequencing is not saturated, therefore we may not be detecting key factors due to technological limitations.

---

**Figure S1.**
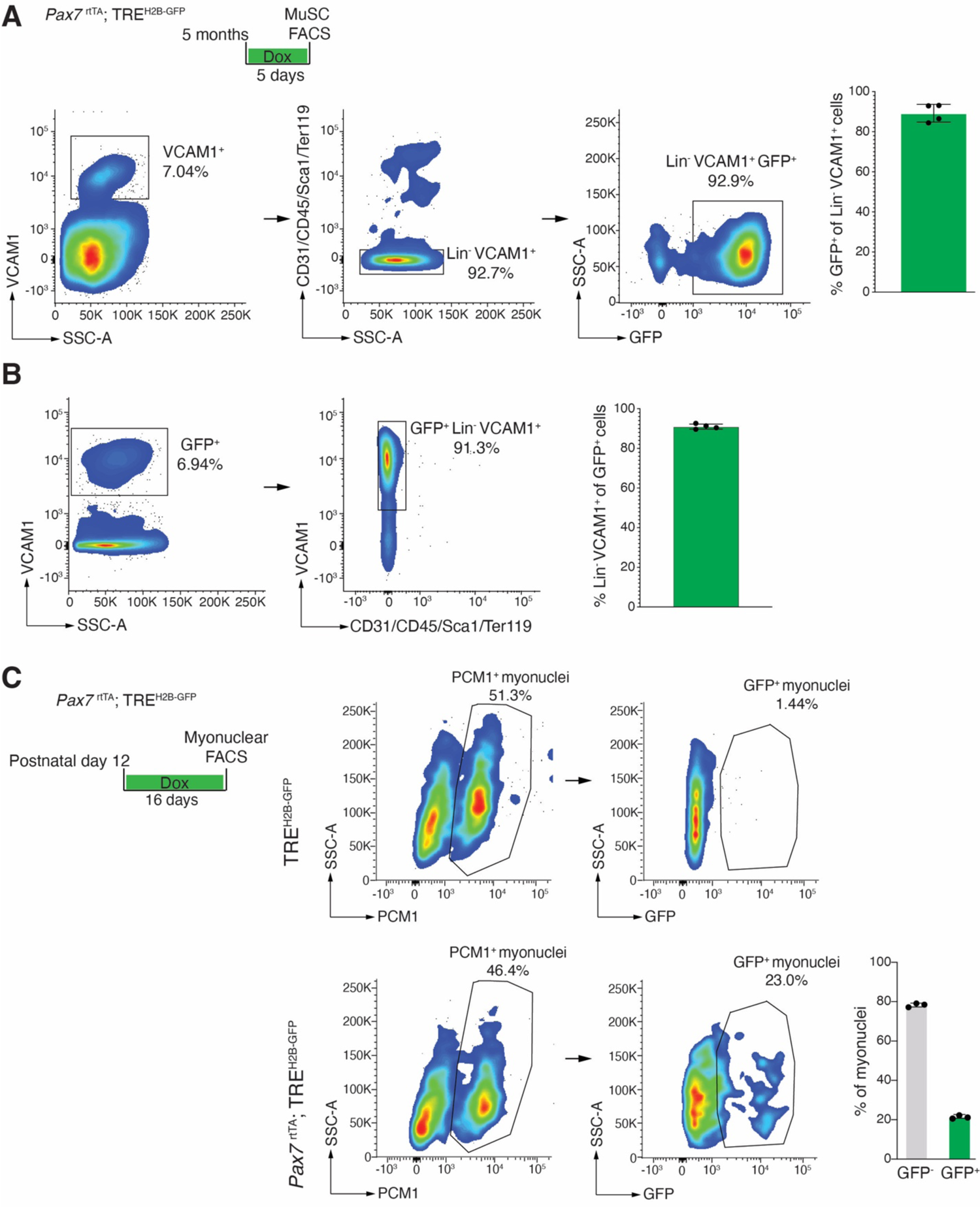
Flow cytometry analysis validates labeling efficiency of MuSCs and myonuclei from *Pax7*^rtTA^; TRE^H2B-GFP^ mice. (A, B) Representative FACS plots for labeling of MuSCs (Lin^-^ VCAM^+^) from *Pax7*^rtTA^; TRE^H2B-GFP^ muscle. Mononuclear cells were digested from gastrocnemius muscles and incubated with antibodies for CD31, CD45, Ter119, Sca1, and VCAM1. (A) Quantification of FACS analysis of CD31^-^CD45^-^Ter119^-^Sca1^-^(Lin^-^) VCAM1^+^ myogenic progenitors expressing GFP. (B) Quantification of FACS analysis to assess GFP expression specificity in CD31^-^CD45^-^Ter119^-^ Sca1^-^(Lin^-^) VCAM1^+^ myogenic progenitors. (n=4). (C) Representative FACS plots for myonuclear labeling (PCM1^+^ GFP^+^) from *Pax7*^rtTA^; TRE^H2B-GFP^ or TRE^H2B-GFP^ mice. Nuclei were isolated from TA muscles and incubated with the PCM1 antibodies. Quantification of FACS plots indicates 20% of myonuclei are labeled with H2B-GFP at P28 after 2 weeks of doxycycline. (n=3). Data are presented as mean ± SD

**Figure S2.**
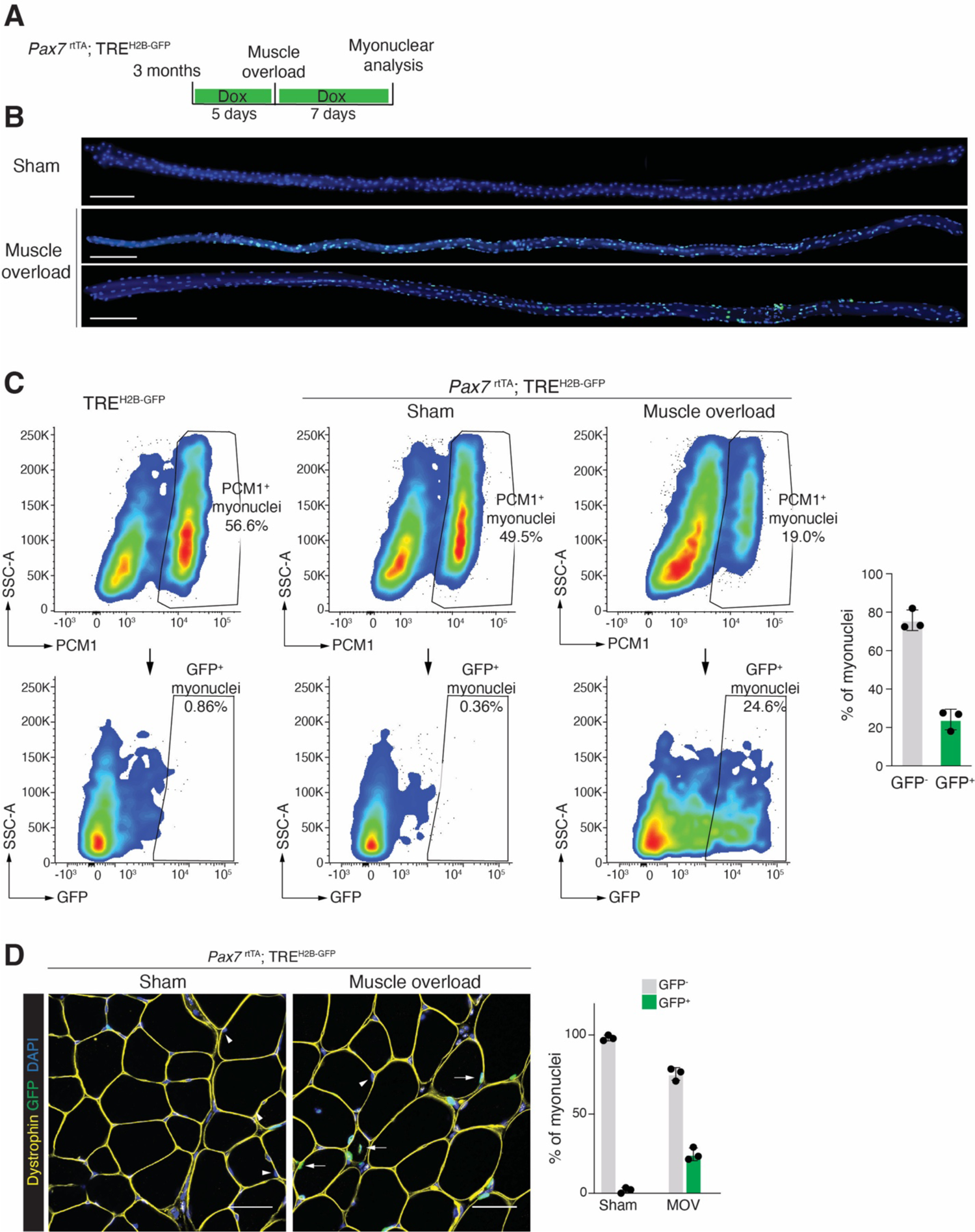
The *Pax7*^rtTA^; TRE^H2B-GFP^ tracking system labels recently fused myonuclei during adult muscle overload. (A) Schematic of the *Pax7*^rtTA^; TRE^H2B-GFP^ lineage tracing system. Doxycycline was administered 5 days before the synergist ablation surgery (muscle overload) for MuSCs labeling. All muscle samples were harvested one week after the surgery. (B) Representative images of single myofibers isolated from the plantaris muscle. Heterogeneous GFP expression within myonuclei was observed in myofibers subjected to muscle overload. Scale bar, 200μm. (C) Representative FACS plot for myonuclear labeling (PCM1^+^ GFP^+^) from *Pax7*^rtTA^; TRE^H2B-GFP^ mice. Nuclei were isolated from plantaris muscle and incubated with the PCM1 antibodies. Quantification of FACS indicates that 20-25% of myonuclei were labeled with H2B-GFP one week after the overload stimulation. (n=3). (D) Representative images showing newly fused GFP^+^ myonuclei (white arrows) and pre-existing GFP^-^ myonuclei (arrowhead) in sham *Pax7*^rtTA^; TRE^H2B-GFP^ mice and after muscle overload (left). Quantification of the percentage of GFP^+^ myonuclei and GFP^-^ myonuclei (right). Scale bar, 20μm. (n=3). Data are presented as mean ± SD.

**Figure S3.**
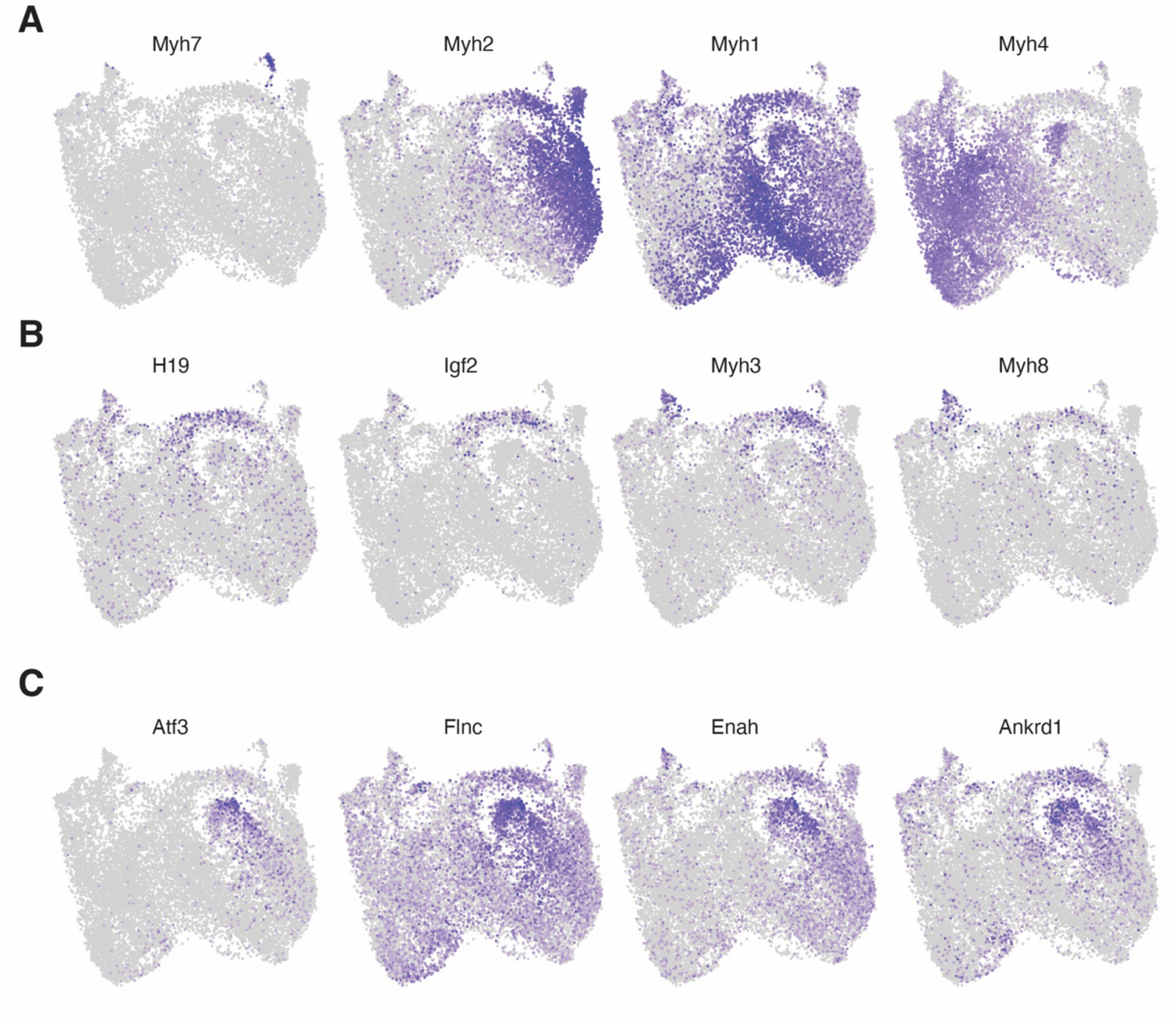
Gene expression in nuclear populations from integrated snRNA-seq data. (A-C) Feature plots of canonical markers of mature myonuclei, and key marker genes of unclassified A, B, and *Atf3*^+^ populations.

**Figure S4.**
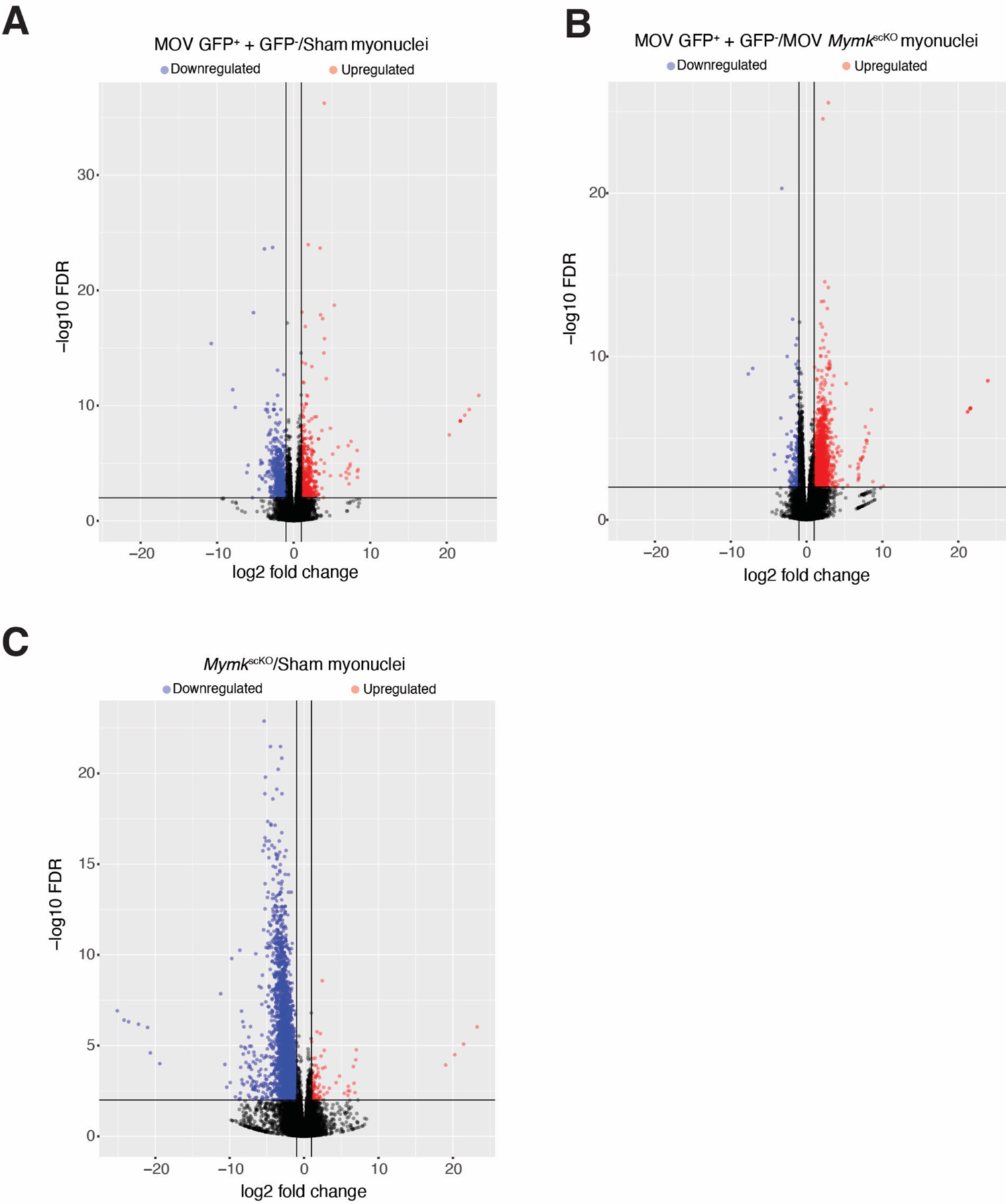
DEG analysis from bnRNA-seq in various myonuclear populations. (A) Volcano plot revealing up- and down-regulated DEGs in WT MOV myonuclei (GFP^+^ and GFP^-^ combined) compared with WT Sham myonuclei. (B) Volcano plot revealing up- and down-regulated DEGs in WT MOV myonuclei (GFP^+^ and GFP^-^ combined) compared with *Mymk*^scKO^ MOV myonuclei. (C) Volcano plot revealing up- and down-regulated DEGs in *Mymk*^scKO^ MOV myonuclei compared with WT Sham myonuclei.

## Methods

### Animals

Animal studies were conducted in accordance with guidelines from the National Institutes of Health and the Division of Veterinary Services (DVS) at Cincinnati Children’s Hospital Medical Center. The IACUC protocol number for this study is #2020-0047. The transgenic mouse lines utilized in this research include *Pax7*^CreER^ ^64^, *Myomaker*^loxP/loxP^ ^46^, *Pax7*^rtTA^, HSA^rtTA^ ^49^, and TRE^H2B-GFP^ ^42^. All mice were maintained on a C57BL/6 genetic background. The *Myomaker*^loxP/loxP^; *Pax7*^CreER^ mice (*Mymk*^scKO^) used in this study were previously described. Tamoxifen (75 mg/kg body weight), dissolved in corn oil with 10% EtOH, was administered to mice through intraperitoneal injection.

The *Pax7*^rtTA^ and TRE^H2B-GFP^ mice were generated by Dr. Christoph Lepper from The Ohio State University. In brief, the targeting construct for homologous recombination in C57BL/6;129sv hybrid ES cells was generated via recombineering method ^65^. A BAC clone (C57BL/6) containing the entire Pax7 locus was obtained from BACPAC Genomics. A chimeric gene cassette consisting of a GPI-anchored YFP fused to HaloTag® (Promega) followed by the reverse tTA (rtTA) gene (Tet-On 3G, Takara Bio), separated via self-cleaving T2A peptide was generated via Gibson Assembly (NEB). This cassette including a 5’ chimeric intron and 3’ poly adenylation signal sequence replaces the sequence of Pax7 exon 1 immediately 3’ of the ATG start codon. For selection in ES cells, a male germline self-excising Ace-Cre-Neo cassette replaces Pax7 exon 2. This Pax7 targeting vector was used for homologous recombination in C57BL/6;129sv hybrid ES cells. Chimeric animals were produced via blastocyst injection of confirmed recombined ES cell clones. Germline transmission of the Pax7rtTA allele was confirmed via genomic PCR. The HSA^rtTA^ mice ^49^ were provided by Dr. John McCarthy from the University of Kentucky. For TRE^H2B-GFP^ expression, mice were given chow supplemented with 0.0625% doxycycline (TestDiet) at the indicated time points.

### Synergistic Ablation (muscle overload)

Plantaris muscles were subjected to muscle overload by implementing bilateral synergistic ablation of the soleus and gastrocnemius muscles, following the procedure described previously^31^. An incision on the lower limb’s posterior-lateral side was made to expose the soleus and gastrocnemius muscles. The distal and proximal tendons of the soleus were cut followed by cutting of the distal gastrocnemius tendon and excision of 75% lateral and medial gastrocnemius.

### Isolation of nuclei

Following euthanasia, tibialis anterior, gastrocnemius, or plantaris muscles were dissected, minced, and placed in homogenization buffer (0.25 M sucrose and 1% BSA in Mg2^+^-free, Ca2^+^-free, RNase-free PBS). An Ultra-Turrax T25 was utilized for further homogenization of the minced tissue. The homogenate was incubated for 5 minutes upon the addition of Triton-X100 (2.5% in RNase-free PBS, added at a 1:6 ratio). All samples were filtered through a 100 μm strainer, centrifuged (3000 × g for 10 minutes at 4 °C) to yield a crude pellet, and resuspended in sorting buffer (2% BSA/RNase-free PBS) before being filtered again via a 40 μm strainer. Nuclei were labeled with Hoechst dye and 0.2 U/μl Protector RNase inhibitor (Roche). The labeled nuclei were purified using FACS (BD Aria, 70 μm nozzle) and gathered in sorting buffer containing Protector RNase inhibitor (0.2 U/μl).

### Immunofluorescence

Muscle sections were fixed using 1% PFA for a duration of 2 minutes, followed by a wash in PBS. The sections were then permeabilized in a solution of 0.2% Triton X-100/PBS for 10 minutes, subsequently washed in PBS, and then blocked with a mixture of 1% BSA, 1% heat-inactivated goat serum, and 0.025% Tween20/PBS for an hour at room temperature in a humidity chamber. All slides were later incubated with rabbit anti-dystrophin primary antibodies (1:100; Abcam) overnight at 4 °C in a humidity chamber. Following the primary antibody incubation, the slides were rinsed in PBS, and then treated with goat anti-rabbit Alexa Fluor 594 secondary antibody (1:100; Invitrogen) for 45 minutes at room temperature in a humidity chamber. Afterward, the slides were washed with PBS and mounted using VectaShield containing DAPI (Vector Laboratories). Upon the completion of immunostaining, images were captured using a Nikon A1R confocal microscope.

### smRNA-FISH on cryosections

Single-molecule FISH assays were conducted using the RNAscope Multiplex Fluorescent Reagent Kit v2 (ACDBio), with slight modifications to the standard protocol. Fresh-frozen cross-sections of tibialis anterior (TA) or plantaris muscles were used as specified. Briefly, 12µm-thick sections were obtained from frozen muscle samples, air-dried for 10 minutes at room temperature (RT), and then immersed in cold 4% paraformaldehyde (PFA) for 20 minutes. The slides were dehydrated sequentially in 50%, 70%, and 100% ethanol for 5 minutes each at RT. Dehydrated slides were preserved in 100% ethanol at −20°C until needed.

All RNAscope analyses on cryosections were co-stained with dystrophin antibodies for analysis of transcriptional signal in myonuclei. To perform immunofluorescence staining, all sections were blocked using a solution containing 1% bovine serum albumin, 1% heat-inactivated goat serum, and 0.025% Tween-20 in phosphate-buffered saline for 30 minutes at RT. The sections were then incubated with anti-dystrophin antibodies (1:100, Abcam) overnight at 4°C, followed by two washes with 1xPBS, 5 minutes each. Subsequently, AlexaFluor secondary antibodies (1:100) (Invitrogen) were applied at room temperature for 1 hour, followed by two washes with 1xPBS, 5 minutes each, and a post-fixation step with 4% PFA for an additional 20 minutes. All slides were then subjected to a 10-minute incubation in hydrogen peroxide at RT, followed by a 20-minute incubation with Protease III at RT. Between each incubation, slides were washed with 1xPBST for 5 minutes. Slides were then hybridized with selected probes at 40°C for 2 hours. The remaining AMP hybridization and HRP incubation steps were performed according to the instructions provided in the ACDBio user manual.

The intensity of the smRNA-FISH signal was assessed using two distinct analytical pipelines in *Pax7*^rtTA^; TRE^H2B-GFP^ or *Mymk*^scKO^ mice. In *Pax7*^rtTA^; TRE^H2B-GFP^, the median smRNA-FISH signal intensity was determined within a specified Region of Interest (ROI), each being a 25 square micrometer circular area encompassing either newly formed myonuclei (GFP^+^) or pre-existing myonuclei (GFP^-^). 10-20 ROIs from each image were included in the analysis. For the analysis of *Mymk*^scKO^ mice and their littermate controls, the median smRNA-FISH signal strength was measured across the entire fiber area within dystrophin-labeled myofibers. NIS Elements software (Nikon) was employed for all signal strength analyses.

### smRNA-FISH on EDL myofibers

Extensor digitorum longus (EDL) or plantaris muscles were harvested and incubated in high-glucose DMEM (HyClone Laboratories) containing 0.2% (EDL muscle) or 0.4% (plantaris muscle) collagenase type I (Sigma-Aldrich) at 37°C in a cell culture incubator. Following a 30-minute incubation, muscles were gently triturated using a wide-bore glass pipette to facilitate myofiber release before returning to the incubator. After a maximum of 1 hour of incubation, muscles were triturated until myofibers detached from the tissue. Isolated myofibers were collected, fixed in 4% PFA/PBS for 20-30 minutes at room temperature, and rinsed in PBS. Myofibers were then arranged on Cell-tak adhesive-coated slides and transferred directly to 100% methanol, stored at −20°C until needed.

For rehydration, slides were washed in a descending methanol/PBS+0.1% Tween-20 (PBST) series (50% MeOH/50% PBST; 30% MeOH/70% PBST; 100% PBST) for 5 minutes each. All slides underwent a 10-minute hydrogen peroxide incubation, followed by a 20-minute Protease III incubation at room temperature. Slides were washed with 1xPBST for 5 minutes between incubations. Selected probes were used to hybridize slides at 40°C overnight. The ACDBio user manual provided guidance for the remaining AMP hybridization and HRP incubation procedures.

### Single nuclear RNA-sequencing (snRNA-seq)

Plantaris muscles from HSA^rtTA^; TRE^H2B-GFP^ mice were used for snRNA-seq in this study. Four pieces of plantaris muscle, pooled from two mice that either received or did not receive synergistic ablation, were used for nuclei isolation using the protocol described above. After filtration via a 40 μm strainer, uuclei were labeled with Hoechst dye and 0.2 U/μl Protector RNase inhibitor (Roche). All GFP^+^ Hoechst stained myonuclei were purified using FACS (BD Aria, 70 μm nozzle) and gathered in sorting buffer containing Protector RNase inhibitor (0.2 U/μl). The nuclei were counted with a hemocytometer, and the concentration was fine-tuned as needed to reach the optimal range for the 10X Chromium chip. The 10X Chromium system was then used to load the nuclei using the Single Cell 3′ Reagent Kit v3.1, following the guidelines provided by the manufacturer. Around 12,000 nuclei were loaded for each operation. Sequencing was performed on an Illumina NovaSeq 6000 System.

### Bulk nuclear RNA-sequencing (bnRNA-seq)

To obtain sufficient material for sequencing, skeletal muscles were pooled together for this analysis. For the early development stage, 4 × Gastrocnemius and 4 × tibialis anterior muscles per replicate were used, while for adult muscle overload, 4 × Plantaris muscles per replicate were used. Nuclei were extracted using the protocol described in the methods section above. The washed nuclei pellets were resuspended in sorting buffer (2% BSA/RNase-free PBS) and labeled with anti-PCM1 (1:500, HPA023374, Sigma-Aldrich) on ice for 45 minutes. Nuclei were washed twice in sorting buffer and then labeled with an Alexa Fluor 647-conjugated secondary antibody (1:100, Invitrogen) at 4°C for 30 minutes, followed by labeling with Hoechst dye. The nuclei were washed again and resuspended in sorting buffer with 0.2 U/μl Protector Rnase inhibitor prior to cell sorting to separate the PCM1-labeled fractions using a FACS cell sorter (BD Aria). The sorted nuclei were then pelleted at 3000 × g for 10 minutes before mRNA isolation using the Direct-zol RNA Microprep kit (Zymo research R2060). All RNA samples were initially amplified using the Ovation RNA-Seq System v2 kit (Tecan Genomics), following the manufacturer’s guidelines. This system amplified RNA samples, producing double-stranded cDNA. Their concentrations were gauged using the Qubit dsDNA BR assay, and the cDNA size of each was determined with an Agilent HS DNA Chip.

For all samples, libraries were constructed using the Illumina’s Nextera XT DNA Sample Preparation Kit. Specifically, 1 ng of cDNA was combined with Tagment DNA Buffer. Tagmentation, which includes fragmentation and adaptor tagging, was executed with the Nextera enzyme, and the samples were then incubated at 55℃ for 10 minutes. NT Buffer was added for neutralization. Libraries were produced via PCR with the Nextera PCR Master Mix and two Nextera Indexes, using the following program: 72℃ for 3 minutes, 98℃ for 30 seconds, 12 cycles of 95℃ for 10 seconds, 55℃ for 30 seconds, 72℃ for 1 minute, and a final cycle 72℃ for 5 minutes. Concentrations of these libraries were measured using the Qubit dsDNA HS assay, and the size was determined using the Agilent HS DNA chip. After pooling the samples, the concentration was optimized to ensure at least 40 million reads per sample. 100 bp paired-end sequencing was performed using a NovaSeq SP (200 cycles) v1.5 flow cell. All libraries were sequenced on the Illumina NovaSeq 6000 system.

### FACS analysis

The labeling efficiency of *Pax7*^rtTA^; TRE^H2B-GFP^ in MuSCs in vivo was analyzed by subjecting harvested gastrocnemius muscles to enzymatic digestion with collagenase and pronase. The resultant mononuclear cell suspensions were treated with antibodies against VCAM1 (Biolegend) for APC detection of myogenic cells, and with CD45, CD31, Sca1, and Ter119 for negative selection (all PE-conjugated, Biolegend). The labeling efficacy of *Pax7*^rtTA^; TRE^H2B-GFP^ in myonuclei in vivo was assessed using nuclei extracted from either TA or plantaris muscle as specified. The identical myonuclei sorting protocol described above was applied for GFP^+^ myonuclei percentage analysis. All FACS analysis was performed on an LSRII platform (BD Biosciences), and the creation of data plots was achieved with FlowJo v10.8.1 software.

### bnRNA-seq data processing

FastQ files were processed using the nf-core/rnaseq v3.11.1 pipeline with minor modifications ^66,67^. Briefly, quality control was performed using FastQC. Low-quality bases and adapter sequences were then trimmed and filtered from the reads using Cutadapt v3.4 ^68^ to improve the overall data quality. Ribosomal RNA sequences were subsequently removed from the aligned data using SortMeRNA v4.3.4 ^69^ to eliminate potential contamination from non-target RNA species. The resulting alignments were sorted and indexed using SAMtools v1.16.1 ^70^, enabling efficient storage and retrieval of the alignment information. Reads were aligned to the mm10 genome obtained from GENCODE (vM21) using STAR v2.7.9a ^71^. For read quantification, we utilized Salmon ^72^, which specializes in estimating transcript abundance from RNA-Seq data. Transcripts were annotated using a modified GTF file obtained from gencode (vM24) containing both intronic and exonic regions for genes. Reads were imported into DESeq2 ^73^ using the tximport command from the tximport package ^74^. All differential expression analysis was performed using DESeq2.

### snRNA-seq data processing

Initial read alignment and quantification of FASTQ files were generated using CellRanger/5.0.1. For each dataset, we corrected for ambient background RNA using cellbender^75^. Reads from cellbender were then imported into Seurat objects ^76^, and nuclei with less than 200 unique features were removed from downstream analysis. Seurat objects with the remaining nuclei then underwent doublet identification using Solo/0.2. Nuclei having higher than the 95^th^ percentile of number of unique features, number of UMIs, and mitochondrial reads were excluded. An additional filtration of any nuclei containing higher than 5% reads mapping to the mitochondrial genome were excluded from all datasets. Datasets were normalized using the SCTransform() command, setting the “vst.flavor” argument to “v2”, and the vars.to.regress argument set to “percent.mt”. Additional regression of cell cycle scores was included in the SCTransofrm <u>vars.to</u>.regress argument only for datasets derived from whole muscle sequencing (P10 and Five Months), as these datasets included non-postmitoic cells. First, each cell was given a score using the CellCycleScoring() function provided by Seurat on the normalized RNA Assay, which was normalized using the NormalizeData() function. using the gene signatures for S Phase and G2M phase provided by Seurat ^77^. These genes were converted to the mouse genome by lowercasing all but the first letter of each gene.

### snRNA-Seq data integration

For each integration, the SelectIntegrationFeatures() and PrepSCTIntegration() were used on a list containing each dataset being integrated ^78^. The identified features output from the previous command are given to the PrepSCTIntegration() command. Then, integration anchors were generated using the FindIntegrationAnchors() command, inputting the identified integration features to the “anchor.features” parameter, and specifying the “normalization.method” parameter as “SCT”. Datasets were then integrated by supplying the anchors to the IntegrateData() command, specifying the “normalization.method” parameter as “SCT”. Dimensionality reduction was performed on all integrated datasets using the FindClusters(), FindNeighbors(), and RunUMAP() commands. In the cases of myogenic subsets, after subset, an additional round of dimensionality reduction was performed using RunPCA(), FindClusters(), FindNeighbors(), and RunUMAP() again. UMAPs were visualized using the DimPlot() function.

### PHATE Trajectory

The postnatal day 10 snRNAseq dataset was subset to contain only myogenic populations from non-specialized compartments. The phate() command provided from the phateR package ^79^ was run on the normalized RNA assay, with the parameter t = 30.

### Gene Expression Visualization

All Heatmaps were created using the DoHeatmap() function provided by Seurat. Feature plots were generated using the FeaturePlot() function. DotPlots were created using the DotPlot() function. Input features for Heatmaps and Dotplots were either manually selected, or generated through the FindAllMarkers() function provided by Seurat. VolcanoPlots for the bnRNA-Seq data were generated using the R package ggplot2 function^80^, plotting log transformed p adjusted values on the y axis and log2 fold change on the x axis.

### Statistical Analyses

We used the following criterion for differentially expressed genes: of log_2_FC≥1, FDR <0.01 (upregulated genes) and log_2_FC≤-1, FDR <0.01 (downregulated genes). Gene ontology analysis was performed using ToppGene (https://toppgene.cchmc.org) ^81^. Sample sizes are noted in the figure legends. Data were processed using GraphPad Prism 9 software. In all graphs, error bars indicated standard deviation (SD). Data were compared between groups using various statistical tests, indicated in the figure legends, and based on number of groups, normality of the data, and variance of standard deviations. The criterion for statistical significance was *p<0.05, **p<0.01, ***p<0.001, ****p<0.0001.

### Data and Code Availability

Raw sequencing data has been deposited in GEO (GSE241035) and are publicly available. Data analysis is described in the Methods. Scripts are available on GitHub: https://github.com/cswoboda/NF-Myonuclei.

## Acknowledgements

We thank members of the Millay laboratory and Vikram Prasad (Molkentin lab) for discussion. We would like to acknowledge the assistance of the Research Flow Cytometry Core, Single Cell Genomics Core, and DNA sequencing and Genotyping Core at Cincinnati Children’s Hospital Medical Center. This work was mainly supported by a grant to D.P.M. from the National Institutes of Health (R01AG059605). Work in the Millay laboratory is also funded by grants to D.P.M. from Children’s Hospital Research Foundation, National Institutes of Health (R01AR068286, R61AR076771), and a sponsored research agreement with Sana Biotechnology. Work for this project in the Weirauch laboratory is funded by grants to M.T.W. from the National Institutes of Health (P30AR070549) and Cincinnati Children’s Hospital Research Foundation (ARC award #53632). The Lepper laboratory is supported by a grant from the National Institutes of Health (R01AR078231).

## Author contributions

C.S., C.O.S., and D.P.M. conceived the project. C.S., C.O.S., M.J.P., S.P., and A.V. conducted experiments and analyzed the data. M.T.W., C.L., and D.P.M. supervised the project. C.S., C.O.S., and D.P.M. wrote the manuscript with input from all authors.

